# Discovery and Quality Analysis of a Comprehensive Set of Structural Variants and Short Tandem Repeats

**DOI:** 10.1101/713198

**Authors:** David Jakubosky, Erin N. Smith, Matteo D’Antonio, Marc Jan Bonder, William W. Young Greenwald, Agnieszka D’Antonio-Chronowska, Hiroko Matsui, i2QTL Consortium, HipSci Consortium, Oliver Stegle, Stephen B. Montgomery, Christopher DeBoever, Kelly A. Frazer

## Abstract

Structural variants (SVs) and short tandem repeats (STRs) are important sources of genetic diversity but are not routinely analyzed in genetic studies because they are difficult to accurately identify and genotype. Because SVs and STRs range in size and type, it is necessary to apply multiple algorithms that incorporate different types of evidence from sequencing data and employ complex filtering strategies to discover a comprehensive set of high-quality and reproducible variants. Here we assembled a set of 719 deep whole genome sequencing (WGS) samples (mean 42x) from 477 distinct individuals which we used to discover and genotype a wide spectrum of SV and STR variants using five algorithms. We used 177 unique pairs of genetic replicates to identify factors that affect variant call reproducibility and developed a systematic filtering strategy to create of one of the most complete and well characterized maps of SVs and STRs to date.

## Introduction

Structural variants (SVs) and short tandem repeats (STRs) respresent a significant fraction of polymorphic bases in the human genome and have been shown to cause monogenic diseases and contribute to complex disease risk (Beck et al., 2015; Brandler et al., 2018; Carvalho and Lupski, 2016; Den Dunnen, 2017; La Spada and Taylor, 2010; Malhotra et al., 2011; Malhotra and Sebat, 2012; McMurray, 2010; Michaelson et al., 2012; Mirkin, 2007; Nelson et al., 2013; Pearson, 2003; Spielmann and Klopocki, 2013; Spielmann and Mundlos, 2013). STRs are polymorphic 1-6 base pair (bp) sequence repeats whose total size can range from ~10bp to more than 1kb while SVs capture diverse sequence variation greater than 50bp in size such as insertions, duplications, deletions, and mobile element insertions (MEIs). The full contribution of STRs and SVs to disease risk, quantitative molecular traits, and other human phenotypes is currently not understood because previous studies have typically genotyped SVs and STRs using arrays or low coverage sequencing which are limited in their ability to accurately identify and genotype these variants in many samples across different variant classes and sizes (Gamazon et al., 2011; Kong et al., 2012; Schlattl et al., 2011; Sudmant et al., 2015). The increasing adoption of high coverage whole genome sequencing data (WGS), however, has recently enabled the development of improved methods to identify STRs and different classes of SVs (Chiang et al., 2017; Hehir-Kwa et al., 2016; Kosugi et al., 2019).

While high-depth WGS data has made it possible to profile a wider spectrum of genetic variation, the variability in the size and characteristics of SV classes necessitates the use of several algorithmic approaches that differ in the types of evidence used to capture all classes of SVs. For instance, some algorithms specialize in identifying small SVs (50-5,000 bp) by using split or discordant read (abnormal insert size) information to determine the location of SV breakpoints with high resolution (Fan et al., 2014; Kronenberg et al., 2015; Layer et al., 2014b; Rausch et al., 2012). Other algorithms detect large SVs (>5 kb) by comparing the amount of reads that align to the reference genome to identify regions that differ in copy number between samples (Abyzov et al., 2011; Handsaker et al., 2015; Klambauer et al., 2012; Zhu et al., 2012), but with lower resolution breakpoint precision (Becker et al., 2018; Chaisson et al., 2019; Hehir-Kwa et al., 2016; Lin et al., 2015). Finally, algorithms have also been designed to contend with more complex multi-allelic signatures, including regions with multiple copy number or repeat alleleles that are more challenging to genotype than biallelic variants (Handsaker et al., 2015; Klambauer et al., 2012). Genotyping SVs and STRs across many samples thus requires using several highly parameterized algorithms to discover each class of SVs, processing schemes to combine results from different algorithms, and detailed filtering to remove false positives or inconsistely genotyped variants. Such pipelines for SV/STR identification must also be sensitive to study-specific parameters such as library prepration methods, sequencing depth, cell/tissue type, and read length (Becker et al., 2018; Chaisson et al., 2019; Chiang et al., 2017; Hehir-Kwa et al., 2016; Kosugi et al., 2019; Lin et al., 2015). Thus, due to the diversity of SV/STR calling algorithms and the need for complex downstream processing, it remains difficult to create a comprehensive SV and STR call set with consistent quality that covers the spectrum of variant sizes and subclasses.

In addition to difficulties associated with complex pipelines for calling SVs and STRs, the need to perform *de novo* discovery and subsequent genotyping of variants across hundreds or thousands of samples leads to inconsistencies between variant calls across studies. A comprehensive catalog of SVs and STRs in the human genome would make it possible for different studies to genotype this same set of variants. While several efforts are underway to establish such catalogs of SVs (Audano et al., 2019; Chaisson et al., 2015; Chaisson et al., 2019; Chiang et al., 2017; Collins et al., 2017; Hehir-Kwa et al., 2016; Levy-Sakin et al., 2019; Sudmant et al., 2015) and STRs (Gymrek et al., 2016; Willems et al., 2017), most are limited in their number and diversity of samples or do not capture all types of variants due to the sequencing depth or algorithms employed. There is also a need to understand the extent to which differences in sample collection and preparation may impact SV and STR calling by measuring the reproducibility of variants called on genetic duplicate samples that share the same genome but were collected and prepped separately. A comprehensive reference catalog of high quality SVs and STRs discovered in a large set of subjects with deep WGS data could therefore be useful for calling and genotyping the full spectrum of variants across future studies involving hundreds to thousands of subjects.

In this study, as part of the i2QTL consortium, we profiled 719 whole genomes from iPSCORE (D’Antonio et al., 2018; DeBoever et al., 2017; Panopoulos et al., 2017b) and HipSci (Kilpinen et al., 2017a; Streeter et al., 2017) with five variant calling algorithms to capture a wide spectrum of of SVs including biallelic deletions and duplications, multi-allelic copy number variants (mCNVs; regions that have more than two copy number alleles segregating in the population), MEIs, reference MEIs (rMEI), inversions, unspecified breakends (BND), and STRs. We identified algorithm-specific quality metrics and SV genomic properties associated with the reproducibility of variant calling using 177 pairs of genetic replicates embedded in our collection (25 monozygotic twin pairs and 152 fibroblast-iPSC pairs) and devised filtering and processing approaches to obtain a highly accurate, non-redundant call set across variant classes and algorithmic approaches. We compared our set of SVs with those identified in GTEx (Chiang et al., 2017) and 1KGP (Sudmant et al., 2015) and found that we capture the vast majority of common SVs likely discoverable in Europeans with short read sequencing and add novel, high-quality variants at lower allele frequencies. Finally, we characterized the extent to which different classes of SVs and STRs are tagged by SNPs and indels. This study establishes methods for filtering SVs and STRs to obtain reproducible variant calls and provides a high-quality reference catalog of SVs and STRs that will benefit studies that investigate how these variants contribute to human disease.

## Results

### The i2QTL sample set

We generated the i2QTL variant calls dataset by calling SNVs, indels, SVs, and STRs using 719 human WGS samples from 477 unique donors. (Figure 1A, Table S1, Table S2). The samples were obtained by combining data from two large induced pluripoitent stem cell (iPSC) resources: 1) iPSCORE (273 individuals, mean WGS coverage 50X, range 36-126X) (D’Antonio et al., 2018; DeBoever et al., 2017; Panopoulos et al., 2017b) and 2) HipSci (446 samples from 204 individuals, mean WGS coverage 37X, range 35-78X) (Kilpinen et al., 2017b; Streeter et al., 2017) (Figure 1B, C). The 477 individuals include members of all five 1KGP superpopulations (Auton et al., 2015): 415 European, 34 East Asian, 15 Admixed American, 7 South Asian, and 6 African. While all 204 HipSci donors were unrelated, there were 183 donors in iPSCORE that are part of 56 unique families (2-14 individuals/family) (Figure S1), including 25 monozygotic (MZ) twin pairs (Figure 1D). For 152 HipSci individuals, we also obtained matched fibroblast and iPSC WGS data (Figure 1D). Between these 152 matched samples and 25 MZ twin pairs, we had WGS data for 177 genetic replicates, which we used to determine quality filtering thresholds and to calculate reproducibility of calls across all variant classes.

**Figure 1.**
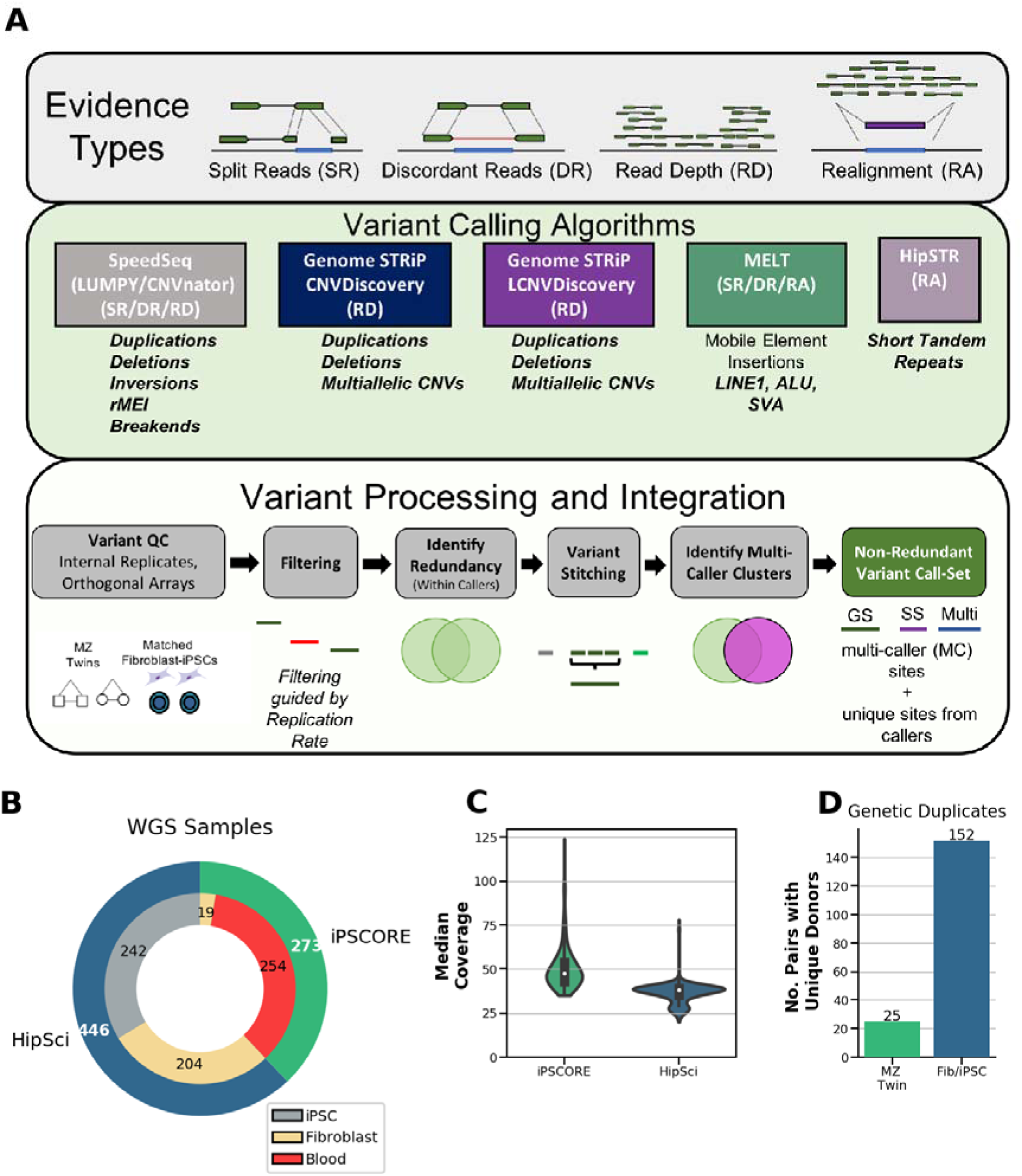
Variant Calling, Processing and i2QTL WGS Samples. (A) Illustration of the evidence types from short read sequencing data utilized in variant calling (top). Description of the variant callers utilized, the types of variants they identify, and the evidence they use (middle). Flowchart showing the processing, quality control (see Methods), and integration of SVs from different variant callers (bottom). (B) Pie chart showing the number of whole genome sequencing samples from the iPSCORE or HipSci studies used for variant calling and the cell type from which DNA was obtained. (C) Distribution of the median coverage of whole genomes from iPSCORE (green) and HipSci (blue). (D) Number of genetic replicate samples included in the collection, including 25 monozygotic twin pairs (iPSCORE) and fibroblast-iPSC pairs from 152 unique donors (HipSci). These data enable robust variant calling for all classes of genetic variation along with reproducibility analysis.

**Table 1.** Summary of i2QTL variants called from samples in the HipSci and iPSCORE collections. Common variants are defined as those with ≥ 5% non-mode allele frequency (NMAF) for SVs and STRs and ≥ 5% MAF for SNVs and indels.

|  | Variant Class | No. Variants | No. Common Variants |
| --- | --- | --- | --- |
|  | SNV | 41,826,418 | 7,013,178 |
|  | INDEL | 7,040,457 | 1,862,365 |
| Copy Number Variants (CNV) | Deletion (DEL) | 16,238 | 3,490 |
|  | Duplication (DUP) | 2,693 | 416 |
|  | Multiallelic CNV (mCNV) | 1,703 | 949 |
|  | Other SV (BND) | 4,612 | 1,377 |
|  | Inversion (INV) | 210 | 92 |
|  | Reference Mobile Element Insertions (rMEI) | 2,343 | 1,689 |
| Mobile Element Insertions (MEI) | ALU | 7,880 | 2,385 |
|  | LINE1 | 1,175 | 262 |
|  | SVA | 442 | 115 |
|  | Short Tandem Repeats (STR) | 588,189 | 381,053 |
|  | **Total SV** | **37,296** | **10,775** |
|  | **Total SV/STR** | **625,485** | **391,828** |
|  | **Total** | **49,492,360** | **9,267,371** |

### Comprehensive structural variant call set

To identify SVs across a wide range of sizes (50 bp to > 1Mb) and classes, we called variants using four algorithms (Figure 1A): SpeedSeq (LUMPY/with CNVnator support)(Abyzov et al., 2011; Chiang et al., 2015; Layer et al., 2014b), Genome STRiP CNVDiscovery, Genome STRiP LCNVDiscovery(Handsaker et al., 2015), and MELT (Gardner et al., 2017). Together, these algorithms incorporate information from two evidence types: 1) read-pair signal (LUMPY and MELT), which includes detection of split reads (two portions of the same read map to different genomic locations) and discordant read pairs (aligned to the genome with abnormal insert size or orientation); and 2) read-depth (Genome STRiP CNVDiscovery, Genome STRiP LCNVDiscovery, CNVnator). Generally, read-pair signal enables discovery of shorter variants (50bp) and balanced events, while read-depth signal is limited to discovery of longer (>1kb) copy number variants (CNVs) which include biallelic deletions, biallelic duplications and multiallelic copy number variants (mCNVs). When variant calling algorithms utilize information from a group of samples to predict genotypes, study-specific differences in the WGS data (cell type assayed, library preparation technique) can cause erroneous variant calls. To account for this, we performed variant calling and genotyping separately in HipSci and iPSCORE samples for Genome STRiP and combined variant calls afterward to avoid batch effects during variant calling (Methods). Using read-pair signals we detected 223,371 SVs consisting of CNvs, inversions, MEIs, and novel adjacencies of indeterminate type referred to as “breakends” (BND). Among these SVs, biallelic deletions and biallelic duplications were also supported by supplementary read-depth evidence (CNVnator). Using read-depth signals alone (Genome STRiP), we detected 28,417 biallelic deletions, biallelic duplications, and mCNVs, bringing the initial call set to a combined 251,788 SVs, before additional processing (Figure S2–S6).

### Reproducibility of SV Calling is Associated with Quality Metrics

Because there is considerable diversity in subtypes of SVs and disparities between detection algorithms, measuring structural variant quality is challenging. Here we used 177 genetic replicates (25 MZ twin pairs and 152 matched fibroblast and iPSC pairs) to measure reproducibility of SV calls for each variant class and SV calling approach under a range of quality metric filter thresholds. Because of complications in variant calling on sex chromosomes due to dosage differences in males and females, we analyzed reproducibility among 198,651 autosomal SVs. Notably, we were able to assess the reproducibility of most variants in the SV call set since 44% of autosomal SVs (88,496) segregated in at least one monozygotic twin pair, 65.4% (129,937) segregated in at least one fibroblast-iPSC pair (Figure 2A), and 71.8% (142,678) segregated in any of the 177 genetic replicates. For each variant that segregated in at least one genetic replicate pair, we assessed reproducibility by calculating how often a non-reference genotype in one replicate pair sample was called concordantly in the other replicate sample, which we define as “replication rate” (RR, Methods). Replication rates were calculated for each SV separately among MZ twin pairs and fibroblast-iPSC pairs. The 25 MZ twin pairs were used to select filters because they have matched cell types and fewer somatic differences(D’Antonio et al., 2018) while the 152 matched fibroblasts-iPSC pairs were used to confirm the performance of these thresholds in the HipSci collection.

**Figure 2.**
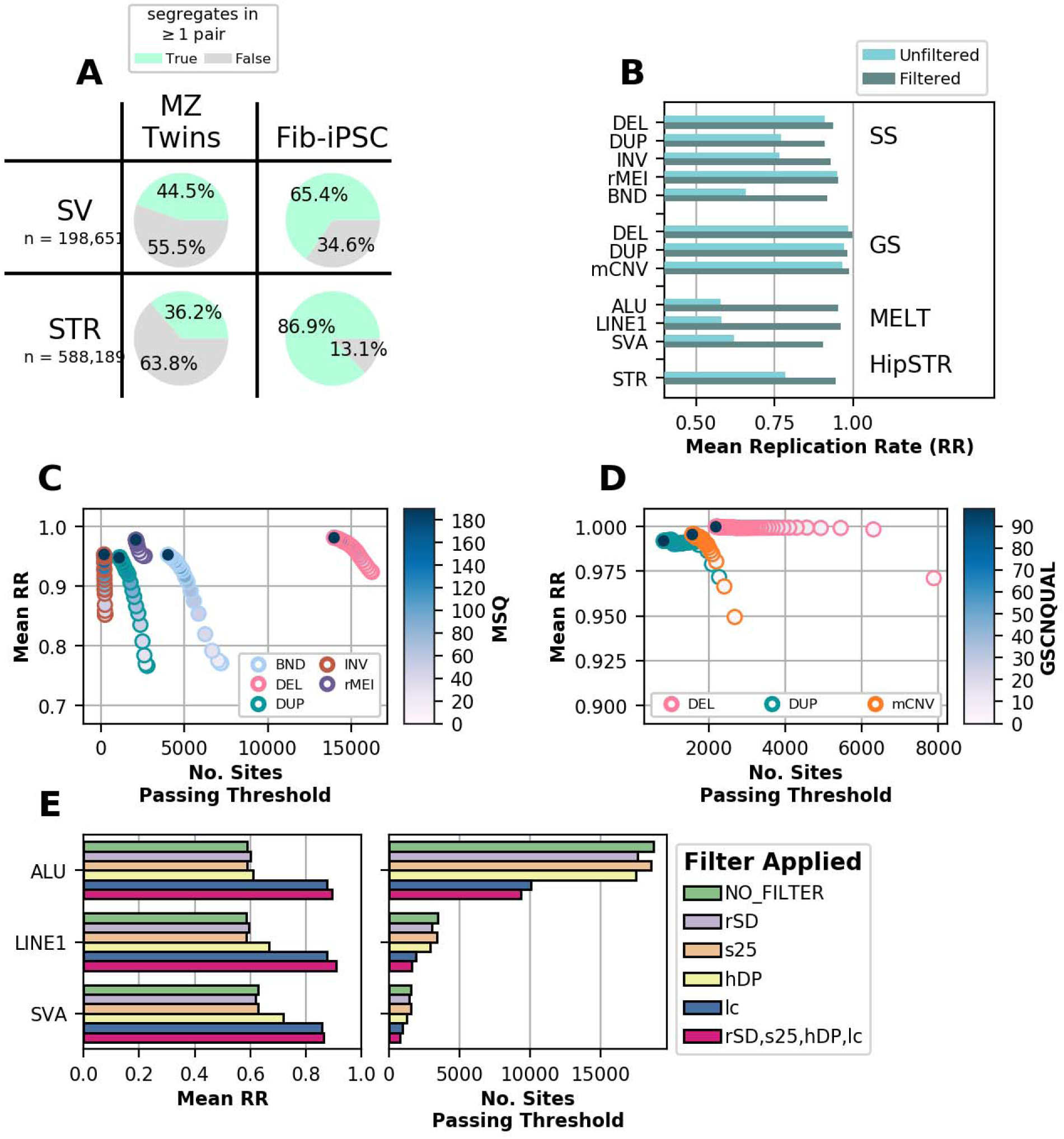
Replication Rate is Associated With Reported Quality Metrics. (A) Proportion of SVs and STRs that were non-reference (green) in at least one of the iPSCORE MZ twin pairs or HipSci fibroblast-iPSC pairs prior to filtering. (B) Replication rate of variants before and after filtering and deduplicating within caller. (C) Replication rate in MZ twins versus the number of total SpeedSeq (LUMPY) sites remaining that pass criteria when filtering variants to different thresholds for MSQ score (indicated by color). (D) Replication rate versus the number of total Genome STRiP sites remaining that pass criteria when filtering variants to different thresholds for GSCNQUAL score (indicated by color). (E) Replication rate in MZ twins for MELT sites that pass criteria when filtering variants under suggested hard site filters (left). Pink represents the result of filtering using all 4 exclusion criteria (rSD, s25, hDP, lc; see Methods). The number of total sites remaining that passed criteria is shown at right.

Prior to filtering on quality metrics (Table S3), we observed that within the 25 MZ twin pairs CNVs (deletions, duplications and mCNVs) detected with Genome STRiP showed high reproducibility (RR > 0.96) as did the SpeedSeq deletions (RR > 0.9) and rMEIs (RR > 0.95), whereas SpeedSeq duplications and inversions (both RR < 0.77), BND (RR=0.65), and MELT MEIs (RR = 0.59) had lower reproducibility (Figure 2B). Interestingly, we found that for all variant callers, increasingly strict quality metric filters yielded variant sets with higher average replication rates, supporting the premise that reproducibility is a predictor of variant quality (Figure 2 C-E). For example, we found strong relationships between Median Sample Quality (MSQ) score from SpeedSeq, the GSCNQUAL score from Genome STRiP, and qualitative filters from MELT and the average RR of filtered variants (Figure 3C-E). Notably, filtering MELT variants called in low complexity regions (“lc” tag in FILTER) improved reproducibility from 59% to 87.5% in MZ twins and applying all four MELT filters improved RR to ~95% (Figure 2E, Methods). Using RR we selected strict quality metric thresholds for each caller and variant class to achieve high specificity without removing a significant number of variants. We observed that within each algorithm, different variant classes required different levels of filtering stringency to attain the same reproducibility (Figure 3C,D). For instance, insertions and duplications were less reliably genotyped than deletions regardless of detection method (Chaisson et al., 2019; Chiang et al., 2017; Kosugi et al., 2019), and SpeedSeq duplications required an MSQ score of 100 to attain >0.9 RR while deletions had an RR of 0.92 with no MSQ filtering (Figure 2C) in MZ twin pairs.

**Figure 3.**
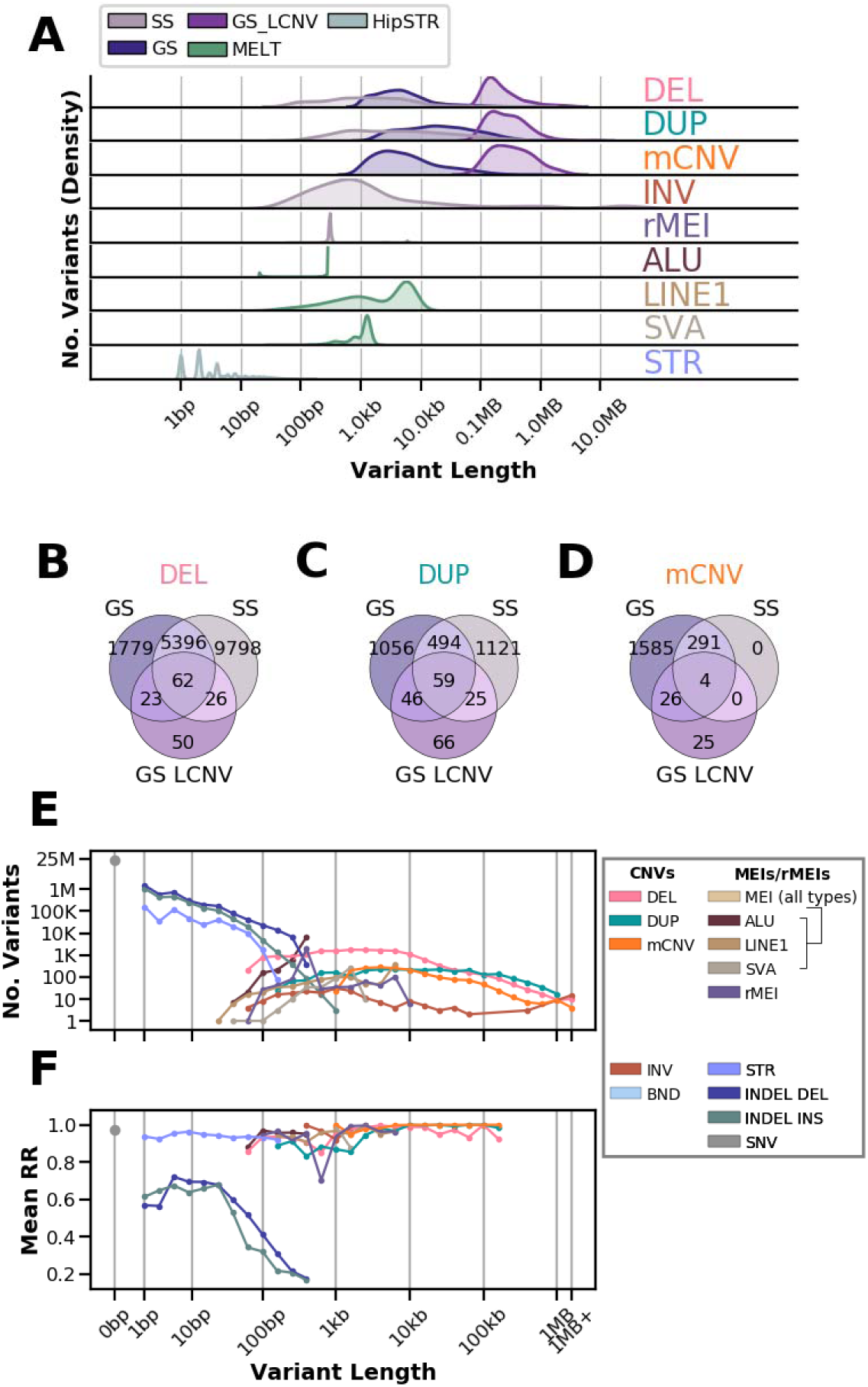
Length Distributions and Intersection between Variants Identified with each Algorithm. (A) Density plot showing the size spectrum of each variant caller before identifying multi-caller clusters. (B-D) Number of overlapping variants after identifying multi-caller clusters for deletions (B) duplications (C) and mCNVs (D). (E) Number of variants in the non-redundant call set separated by variant class and grouped in log linear bins by variant length. Points are drawn at the upper limit of each bin (eg. a bin from 50-100bp is drawn at 100bp). For STRs length represents the maximum number of bases different from the reference at each site (largest insertion or deletion observed). (F) The average replication rate of variants segregating in the 25 monozygotic twin pairs is represented for each length bin that contains at least 10 variants. GATK SNVs and indels previously discovered in iPSCORE samples (DeBoever et al., 2017) were used for (E) and (F).

After filtering, we obtained 50,980 autosomal variants (20.2% of initial call set) with generally high RR (>0.9) for all callers, although variants called by SpeedSeq and MELT tended to have lower RR than those called by Genome STRiP (Figure 2B) suggesting that variants called using read pair signal are less reproducibly genotyped between genetic replicates than than those called by read depth signals. We tested for batch effects by comparing allele frequencies between iPSCORE and HipSci samples and found that they largely agreed across algorithms (Figures S2, S3, S6). We compared the CNV genotypes to those called from SNP arrays for 216 iPSCORE samples and found that the FDR for CNVs ranged from 3-7.8% depending on the SV type and algorithm, consistent with previous reports (Chiang et al., 2017; Sudmant et al., 2015). We also found that biallelic SVs generally obeyed Hardy-Weinberg across algorithms after filtering (Figures S2, S3, S6). Together, these results suggest that our stringent filtering approach can be used to obtain comparable, high quality variants across SV classes and algorithms.

### Creating a High Quality, Non-Redundant SV Call Set

SV calling algorithms overlap in the types and sizes of variants they identify (Figure 3A) which can lead to the same genetic variant being called with slightly different breakpoints by different algorithms in the same subject or by the same algorithm in different subjects. To obtain a non-redundant map of structural variation, we devised a graph-based approach to consolidate overlapping sites that are redundant with each other (Figure S7, Methods). We first clustered overlapping variants that were detected using the same algorithm and showed high genotype correlation and designated each cluster as a single distinct SV with a breakpoint defined by the highest quality variant (Figure 1A). We next stitched together neighboring variants from Genome STRiP whose genotypes were correlated because they likely represent a single variant that Genome STRiP called as multiple adjacent variants (Chiang et al., 2017). Finally we clustered overlapping variants identified by different algorithms with high genotype correlation and designated each “multi-caller” cluster as a single distinct SV (Figure 3B-D, Methods). We inspected variants identified by multiple algorithms and found that overlap between Genome STRiP and SpeedSeq was highest among deletions (55%), while duplications and mCNVs were only co-discovered 17% and 15% of the time respectively reflecting both the different size spectrums captured by the two methods (SpeedSeq captures smaller variants) and that evidence types (read-pair/read depth) do not always co-occur. SVs identified by more than one algorithm (i.e. with support from both read pair and read depth signals) had higher replication rates than SVs detected with a single algorithm (Figure S8), supporting the premise that the highest quality sites also tend to be the most reproducible. Overall, we collapsed 50,980 variants to 37,296 non-redundant SVs which were used for downstream analyses (Table 1, Table S4, Figure S9,S10). We examined the numbers and proportions of non-reference calls for each of the 719 i2QTL samples (from 477 individuals) across variant calling algorithms and variant classes (Figures S2-3, 5-6). We observed high consistency in the number of variants per sample except for individuals with African ancestry who had more SVs per sample, consistent with other variant types (Ramachandran et al., 2005; Sudmant et al., 2015). Taken together, these results show that the set of i2QTL SVs is of high quality and demonstrates the utility of using genetic replicate samples for SV filtering and processing.

### Variant Length, Allele Frequency and Reproducibility

Since SVs can vary widely in size and we are using short read data to call SVs, we assessed whether replication rate was related to SV length. While we could detect many more short SVs (< 1kb) than long SVs, we observed that long SVs had higher RR (Figure 3E, F). Generally, SVs greater than 1kb were highly reproducible (> 95% RR) while shorter duplications and insertions tended to have the lowest RR, reflecting the relative lack of consistency in genotyping small read-pair based SVs. This dependence on length was observed across variant calling approaches and independent of allele frequency (Figure S11A,C). We also found that rare variants were slightly less reproducible than common variants across SV classes (Figure S11B,D). These results highlight that it remains challenging to identify SVs in intermediate size ranges (~200 bp to 1 kb) using short read sequencing, because the interval is: 1) too small to distinguish from “noise” in read depth signal; 2) within the bounds of variability in insert size, making discordant read-signal undetectable; and 3) too long to be directly sequenced with a single read. While challenges in the discovery of SVs in the ~200 bp to 1 kb range still exist, the i2QTL call set consists of high-quality SVs across a wide size range of SVs (~50 bp to >1 Mb).

### Comparison between SVs in i2QTL, GTEx and 1KGP

We next investigated what proportion of the SVs in the i2QTL call set are novel compared to previous SV call sets by comparing the the 37,296 non-redundant i2QTL SVs with the existing 1KGP (Sudmant et al., 2015) and GTEx (Chiang et al., 2017) SV call sets. GTEx used 148 deeply sequenced genomes and the 1KGP project used 2,504 shallowly sequenced genomes (7.4x) to call the same SV classes present in i2QTL (excluding BNDs in 1KGP and nonreference MEIs in GTEx) and are therefore strong benchmark datasets. The i2QTL SV call set captured the vast majority of common deletions, duplications, mCNVs, inversions, rMEIs and MEIs present with non-mode allele frequency (NMAF) greater than 0.05 in either study, including 77% of variants present in 1KGP Europeans and 79% of variants present in GTEx (Figure 4A,B). Out of all SV classes, we captured the smallest proportion of common GTEx duplications (49%) and BNDs (17%), likely due to differences in filtering stringency, WGS data quality, and breakpoint merging approaches. In total, 83% of common i2QTL SVs (NMAF > 0.05) were co-discovered by one or both of these studies (Figure 4C). Common deletions had the highest co-discovery rate (87%) while mCNVs had the lowest (~66%), consistent with the idea that mCNV discovery benefits from high read-depth and large numbers of samples (Handsaker et al., 2015). Rare variants (NMAF < 0.05) were more likely to be unique to either set, with ~40% of sites from GTEx and 1KGP represented in the i2QTL call set (Figure 4C). In total, 43% of i2QTL SVs were not found in either GTEx or 1KGP. These novel variants were predominantly rare, tended to have shorter lengths, and, excluding those identified by Genome STRiP, had on average 12% lower replication rates than co-discovered variants (Figure 4C, S12–S14). This is expected given that small SVs are the most difficult to genotype and rare variants are more likely to be false positives or negatives (Figure S11). Overall the i2QTL call set contains a significant number of novel, high-quality variants at lower allele frequencies missing from 1KGP and GTEx and captures most common SVs present in 1KGP and GTEx indicating that the call set contains most common SVs in Europeans.

**Figure 4.**
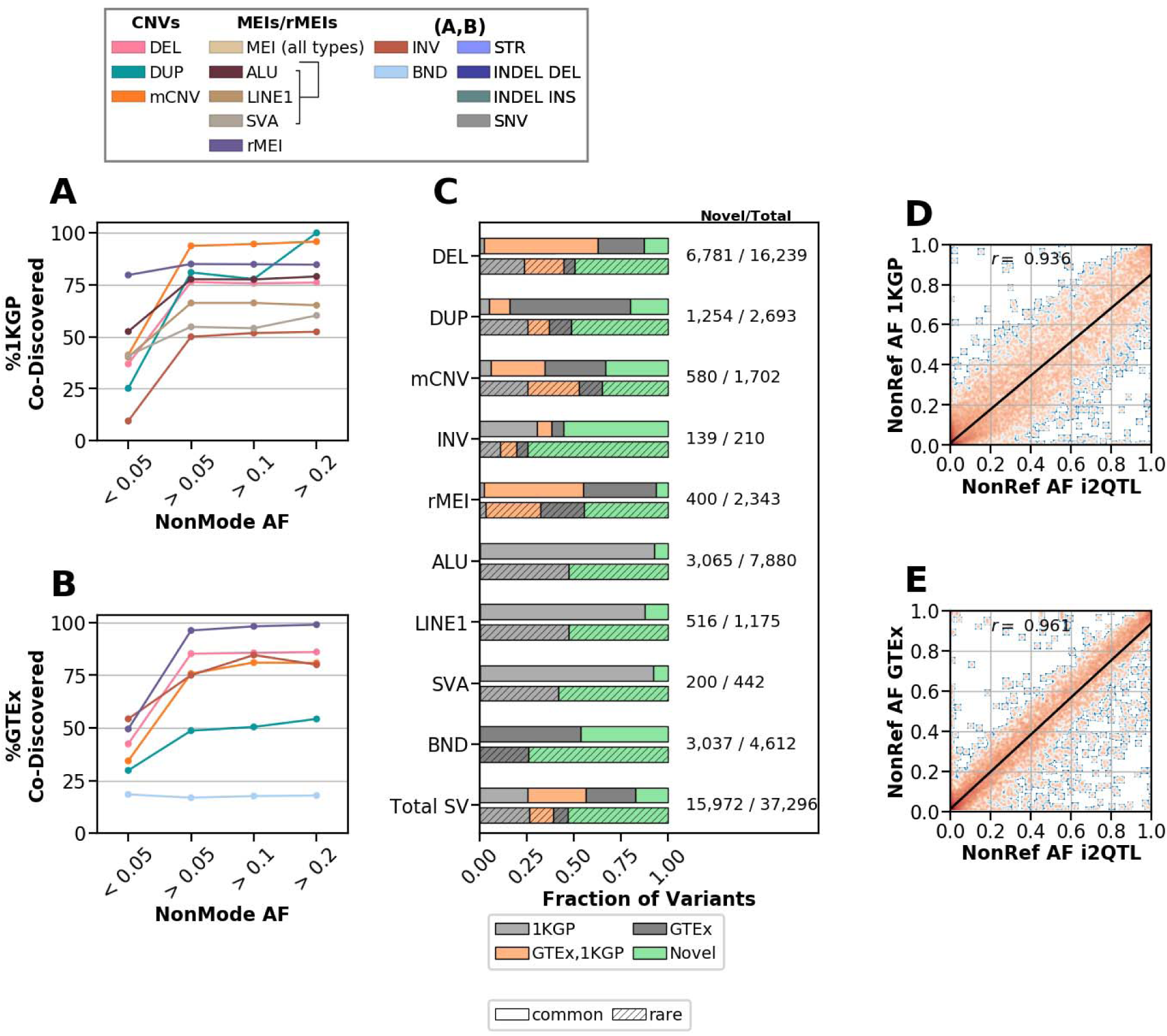
Comparison to other SV calling studies. (A,B) The fraction of variants from either (A) 1KGP (European population) or (B) GTEx that were also captured in our study in different non-mode allele frequency (NMAF) bins. (C) Fraction of i2QTL SVs that were co-discovered in 1KGP, GTEx, both 1KGP and GTEx, or were unique to i2QTL (novel), divided by whether variants were common (> 0.05 NMAF) or rare (< 0.05 NMAF) in unrelated i2QTL samples indicated by absence or presence of hatching respectively. (D,E) Non-reference allele frequency of variants co-discovered in i2QTL and (D) 1KGP (Europeans) or (E) GTEx in their respective discovery samples. Here, the non-reference allele frequency among unrelated i2QTL donors is used, and the density is plotted with orange indicating more observations, and blue fewer.

To assess how similar genotyping sensitivity was between studies, and confirm that overlapping sites were likely to have the same breakpoint, we compared the non-reference allele frequencies of sites that we classified as co-discovered. We found that overall the non-reference allele frequencies of i2QTL variants were highly correlated (r >0.9) with their matched GTEx and 1KGP variants (Figure 4D,E). This was true across variant classes in both studies, with the exception of duplications in 1KGP, which were less correlated (r=0.74), likely as a result of limited genotyping sensitivity in 1KGP due to the use of low coverage WGS data (Figure S15,S16). These results support that the i2QTL SV call set is accurate and contains most common SVs discoverable using short read sequencing data as well as novel, rare SVs, making it a valuable resource for examining functional differences between the SV classes (Jakubosky et al., 2019).

### STR genotyping

We genotyped STR variants at over 1.6 million reference sites using HipSTR (Willems et al., 2017) which employs a hidden Markov model to realign reads around each STR locus (Figure 1A). HipSTR models PCR stutter artifacts to genotype STRs and because of such artifacts, greater genotyping sensitivity and accuracy of predicted *de-novo* STR alleles can be achieved with PCR-free WGS data. In light of this, HipSci samples, which were generated with a PCR-free library preparation, were genotyped separately and then these alleles were used as a reference to genotype iPSCORE samples, which were prepared using a PCR-based library prep, and the results for both sample sets were combined into one call set with consistent alleles. To retain only high-quality STR calls, we applied the genotype specific filters suggested by HipSTR (Willems et al., 2017) and required all sites to have an 80% call rate in iPSCORE or HipSci samples. This resulted in 588,189 autosomal variants with high reproducibility across the range of gentotyped expansion/deletion sizes (1 bp to 150 bp) (overall 94.5%, > 90% in all size bins); these variants were substantially more reproducible than indels in this same size range called by GATK in the i2QTL call set, which overall showed low replication rates (62%) (Figure 3E-F, Figure S17–18). Because HipSTR STRs and GATK indels overlap in size and location, it is likely that some variants are present in both datasets. To compare the genotyping quality of these possibly redundant variants, we intersected GATK indels with 1.6 million HipSTR STR reference loci (Figure S11E). Interestingly, we found that indels (2-100 bp) called by GATK that overlapped an STR locus that was genotyped non-reference in at least one sample by HipSTR had higher RR (77.3%) than those that overlapped STR loci not genotyped as polymorphic by HipSTR (56%), or those that did not overlap an STR region (64.7%). These findings suggest that it is useful to filter large GATK indels (>30bp) because they have low RR (42%), and that STR genotypes are more reproducible than GATK indels.

### Linkage disequilibrium tagging for SVs and STRs

Given the large amount of GWAS and QTL studies performed using genotyping arrays, we next asked to what extent different classes of SVs and STRs are tagged by SNPs and indels. We calculated the maximum linkage disequilibrium (LD) between common SVs and STRs (NMAF > 0.05) within 1Mb of an expressed gene in iPSCs and SNPs/indels (Methods) (Jakubosky et al., 2019) within 50kb of the SVs/STRs. We found that 97.7% of STRs are tagged by a SNP or indel with R^2^ > 0.8 while SVs classes ranged from 44.2% to 86.7% of variants tagged with R^2^ > 0.8 (Figure 5). Duplications and mCNVs were the most poorly tagged classes likely because they are often located near segmental duplications where SNPs and indels are poorly genotyped (Chaisson et al., 2015; Handsaker et al., 2015; Sudmant et al., 2015). These results indicate that most common STRs and some classes of SVs are assayed well by proxy using SNP and indel genotypes, but to increase the coverage of SVs, particularly mCNVs and duplications, studies need to include the genotyping of these variant classes in their samples.

**Figure 5.**
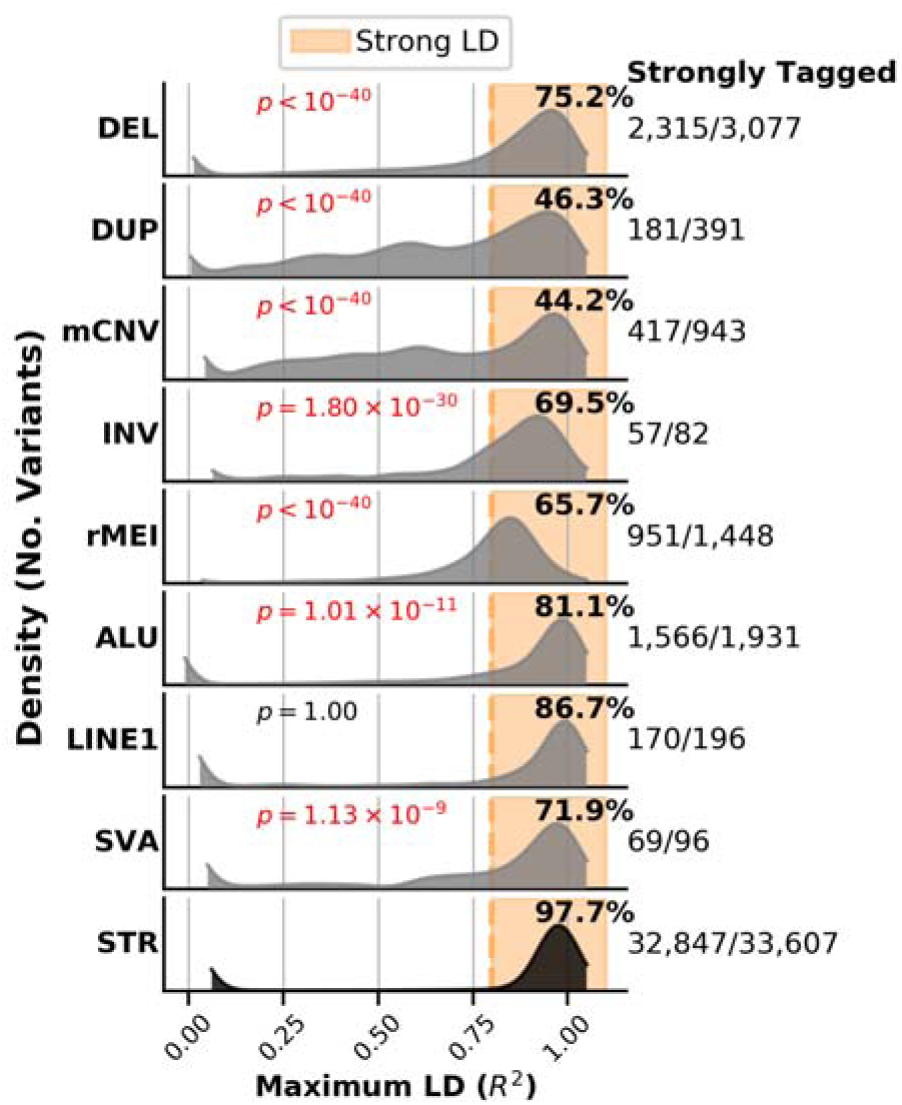
Linkage Disequilibrium of Structural Variants and Short Tandem Repeats with nearby SNPs and Indels. Distribution of maximum linkage disequilibrium (*R^2^*) between common SVs and STRs (non-mode allele frequency > 0.05) and SNVs or indels within 50kb, considering only SVs/STRs that are within 1MB of an expressed gene in iPSCs.

## Discussion

In this study, we discovered and genotyped SVs and STRs in 719 high-coverage WGS samples from 477 unique donors. We detected a wide spectrum of variants across different sizes as most STRs are in the 10bp to 1kb range whereas SVs may span more than 100 kb. We leveraged genetic replicates, such as twin pairs and fibroblast-iPSC matched samples, to test variant calling accuracy and determine filtering approaches to retain only high-quality SVs and STRs. Our filtered call set has very high replication rate (on average >90% for all SV callers), indicating high genotype quality for detected SVs and STRs. The call set captures most of the common variants identified in 1KGP (Sudmant et al., 2015) and GTEx SV variant calling efforts and also contributes novel short (~100-1,000 bp) and rare (NMAF < 5%) variants. The high quality, non-redundant i2QTL SV set described here will serve as a useful reference for other studies and is particularly valuable for genetic association analyses that aim to identify SVs that influence disease risk or quantitative molecular traits like gene expression.

We used five algorithms designed for calling variants across many samples to detect different classes of SVs and STRs and compared the RR in genetic replicates (MZ twin pairs and fibroblast-iPSC pairs) to identify factors that impact RR. We found that we needed to call variants separately in the iPSCORE and HipSci WGS collections and implement specific filtering strategies to account for dataset-specific features such as library preparation techniques to achieve high RR. Given the variability in library preparation methods, future improvements to SV calling algorithms may explicitly adjust for specific library features such as PCR-free sequencing. We also observed differences in RR between different classes and sizes of SVs and different algorithms. We found that SVs in the 100-1,000bp range remain harder to identify and genotype likely due to the limitation of using short reads. We also observed that accuracy was highest for large CNVs (>10 kb) duplications, deletions, and mCNVs suggesting that FDR estimates from orthogonal data sets such as arrays may overestimate accuracy for SV call sets since they generally assess the largest and easiest-to-genotype variants. Future studies that combine deep short read WGS with long read sequencing data may be able to improve the detection and genotyping of SVs in the 100-1,000bp range by directly sequencing them or assembling the short and long reads.

We used genetic replicates to identify algorithm- and SV-specific thresholds and applied these thresholds to filter the initial set of SV calls and create a high quality catalog of SVs and STRs that complements previous SVs identified using low depth WGS or fewer samples (Chiang et al., 2017; Sudmant et al., 2015). We also developed approaches for collapsing redundant SVs and harmonizing SVs called by different algorithms across hundreds of samples. Comparing our SV catalog to previous sets of SVs from the 1KGP and GTEx projects shows that the i2QTL SV call set captures most common (NMAF > 0.05%) SVs in Europeans. However, consistent with others types of genetic variants, we found that African ancestry samples had more SVs than Europeans. Future sequencing studies are needed to fully catalog SVs in other ancestries and identify rare, population-specific SVs. Such multi-ancestry SV catalogs will be indispensable for population sequencing studies such as All of Us (Sankar and Parker, 2017) that aim to integrate genetic and health data for patients from diverse and admixed ancestries.

The filtering scheme and catalog of SVs and STRs presented here can be used in future genetic association and sequencing studies that aim to study the impact of SVs/STRs. One method for utilizing this catalog for calling SVs and STRs is to impute variants via tagging SNPs and indels; a benefit of this approach is that imputation is possible using both array- and sequenced-based genotyping. A second option when sequencing data is available is to skip the *de novo* SV and STR discovery step and instead genotype the reproducible variants reported here. This will restrict genotyping to high-quality sites and may lessen the burden of filtering variant calls. A third option is to perform *de novo* discovery, genotyping, processing, and filtering using the approaches and thresholds that we have identified. While it may be possible that some filtering thresholds need to be adjusted for specific studies, the thresholds provided here likely provide a good starting point for genotyping and filtering *de novo* discovered SVs and STRs in other datasets.

Overall, this study provides a roadmap for discovering and genotyping SVs from WGS data and establishes a high-quality catalog of SVs and STRs that can be used in future genotyping efforts. A companion paper (Jakubosky et al., 2019) examines how the i2QTL SVs and STRs characterized here influence gene expression and contribute to disease risk. These studies demonstrate that SVs and STrs can be reliably identified and genotyped for hundreds of samples and used to study the impact of this class of genetic variation on human health.

## Methods

### Abbreviations

1KGP: 1000 Genomes Project

Indel: Small insertion/deletion variant

SV: Structural Variant

SNV: Single nucleotide variant

SNP: Single nucleotide-polymorphism

WGS: Whole-genome sequencing

FDR: False discovery rate

MAF: Minor Allele Frequency

NMAF: Non-Mode Allele Frequency

MSQ: Median Sample Quality

### Variant Callers

- SS: SpeedSeq SV pipeline (LUMPY read-pair evidence with read depth support from CNVnator)
- GS: Genome STRiP CNVDiscovery pipeline (read depth evidence),
- GS LCNV: Genome STRiP LCNVDiscovery pipeline (read depth evidence)
- MELT: MELT mobile element insertion discovery
- HipSTR: HipSTR short tandem repeat genotyper

### Types of Genetic Variants Detected

- DEL: Biallelic deletion ascertained by LUMPY, GS, GS LCNV
- DUP: Biallelic duplication ascertained by LUMPY, GS, GS LCNV
- mCNV: multiallelic copy number variant ascertained by LUMPY, GS, GS LCNV. This is defined as a variant that has at least 3 predicted alleles.
- INV: inversion ascertained by LUMPY
- rMEI: reference mobile element insertion
- BND: generic “breakend” ascertained by LUMPY. May include deletions and duplications that lack read-depth evidence, balanced rearrangements (INV), MEI or other uncategorized break points.
- ALU: Non-reference Alu element insertion identified by MELT
- LINE1: Non-reference LINE-1 element insertion identified by MELT
- SVA: Non-reference SVA (SINE-R/VNTR/Alu) element insertion identified by MELT
- STR: short tandem repeat variant, detected by HipSTR. Included variants have at least one individual with a change in length from the reference.
- CNV: copy number variant (deletion or duplication structural variant). Encompasses DEL, DUP, mCNV
- MEI: Non-reference mobile element insertion ascertained by MELT, including ALU, LINE1, and SVA elements

## 1. Subject enrollment

273 subjects were recruited as part of the iPSCORE study, of which 215 subjects have been previously described (D’Antonio et al., 2018; DeBoever et al., 2017; Panopoulos et al., 2017b). Data for additional 204 subjects was obtained from the HipSci Collection(Kilpinen et al., 2017a; Streeter et al., 2017). The iPSCORE collection was approved by the Institutional Review Board of the University of California at San Diego (Project #110776ZF). Each of the subjects provided consent, filled out a questionnaire, had blood drawn, and had a 1 mm skin biopsy taken from which fibroblasts were obtained. Five individuals provided consent only for cardiovascular studies, therefore they were removed from downstream analyses. Family relatedness, sex, age, and ethnicity were recorded in the questionnaire. Detailed pedigree information for iPSCORE available in Panopoulos et al. (D’Antonio et al., 2018; DeBoever et al., 2017; Panopoulos et al., 2017b) (dbGAP: phs001035). In total, we utilized a total of 477 HipSci and iPSCORE subjects, 276 were females and 201 were males, and collectively subjects ranged in age from 5 and 89 years of age (Figure S1A). Notably, iPSCORE individuals were included in 56 families composed of two or more subjects (range: 2 to 14 subjects) and 86 single individuals (Figure S1B, Table S1). Overall, 167 iPSCORE individuals were unrelated. All iPSCORE individuals were grouped into one of five superpopulations (European, African, Admixed American, East Asian, and South Asian) on the basis of genotype data (D’Antonio et al., 2018; DeBoever et al., 2017; Panopoulos et al., 2017b) and HipSci samples were similarly categorized (Kilpinen et al., 2017a) (Figure S1C). For HipSci, some subjects had multiple iPSC clones with WGS. For these subjects, we chose the pair of fibroblast and iPSC WGS samples that had the highest reproducibility for Genome STRiP calls (see Section 3.3 below).

## 2. WGS Processing

*IPSCORE:* WGS sequencing for iPSCORE individuals has previously been described in detail DeBoever et al. (DeBoever et al., 2017) and is available on dbGaP (dbGAP: phs001035). DNA isolated from either blood (254 samples) or fibroblasts (19 samples) (Table S2, Figure 1A), was PCR-amplified and sequenced on Illumina HiSeqX (150 base paired-end). We obtained an average of 180.9 billion total raw bases per sample (range 117.81 to 523.49 billion bases). The quality of raw fastq files was assessed using FASTQC (Brown et al., 2017). Reads were then aligned to the human b37 genome assembly with decoy sequences included and a Sendai virus contig with the BWA-mem algorithm under default parameters (Li and Durbin, 2009).

*HipSci:* We downloaded cram files associated with 446 genomes (mean depth 36.3X) generated with a PCR free protocol from 204 healthy donors (ENA Study Accession: ERA828) (Kilpinen et al., 2017a). Genomes were previously aligned to hs37d5 genome, a reference identical to the one used for iPSCORE alignments with the exception of the inclusion of a Sendai virus contig. Cram files were converted to the bam file format and merged when necessary using samtools(Li et al., 2009) Bam files from both iPSCORE and HipSci were sorted with sambamba (Tarasov et al., 2015) and duplicates were marked with biobambam2 (https://gitlab.com/german.tischler/biobambam2).

## 3. Variant Calling, Processing, and QC analysis of SV and STRs

### Code Availability

Code used for analyses and variant processing can be found on GitHub (https://github.com/frazer-lab/i2QTL-SV-STR-analysis).

### 3.1 Quality Control Methods Overview

#### 3.1.1 Replication rate and filtering strategy for SVs and STRs

To minimize the number of poorly genotyped structural variants (SVs) and maximize quality across multiple variant calling approaches, we used the replication rate (RR) metric, calculated as the proportion of non-reference genotypes that were also called non-reference in a paired genetic replicate, as a measure of the reproducibility (and quality) of a variant. The rationale behind this approach is that variants that have high genotyping accuracy should be genotyped consistently in different samples with the same genome and that variants with low genotyping accuracy will differ between samples with the same genome. Under this logic, variants should be consistently genotyped in samples with the same genomes (e.g. technical duplicates, monozygotic twins) and discrepancies would result from false negative or false positive genotypes.

To determine RR for all variant classes, we used genetic duplicate samples in the form of monozygotic twin pairs (n=25) and fibroblast iPSC pairs (n = 152). We used RR to assess the reproducibility of variants under different filtering conditions; the filters were specific to the unique quality metrics measured by each calling algorithm. Using the relationships between filters and RR that we identified, we selected filtering criteria for each variant class in each caller to maximize the quality (specificity) and the number of variants (sensitivity) called.

Because there may be a greater number of somatic variations between fibroblasts and iPSC clones (D’Antonio et al., 2018) due to reprogramming, replication rates in monozygotic twins were used to select thresholds, and iPSC-fibroblast pairs were used for additional confirmation. For this analysis, one member of each pair of genetic duplicates was chosen arbitrarily as the “comparison sample”, and the concordance of non-reference sites in this sample was assessed with respect to the other sample. Replication rate was calculated on all autosomal SVs on a site-by-site basis as the number of pairs with matching non-reference genotypes divided by the total number of pairs with at least one non-reference genotype. Average RRs reported for particular SV classes were calculated as the average RR over all SVs in that class.

#### 3.1.2 Batch effects and Hardy Weinberg Equilibrium

The i2QTL Consortium includes WGS data from iPSCORE and HipSci (Kilpinen et al., 2017a; Panopoulos et al., 2017a), which are different in aspects which may affect variant calling: 1) mean coverage is higher for iPSCORE (50.4X, compared with 36.6X); 2) while most iPSCORE donors had WGS from blood and only 14 from skin fibroblasts, all HipSci donors had WGS from skin fibroblasts; and 3) HipSci samples were sequenced using a PCR-free protocol (Figure S1, Figure 1, Table S2). To limit the batch effects associated with these differences, in cases where a variant caller used information from the entire set of samples to build a global model (Genome STRiP (Handsaker et al., 2015) and HipSTR (Willems et al., 2017)), we genotyped or performed discovery separately in iPSCORE samples and HipSci samples, which were additionally divided into two groups for fibroblast and iPSC samples.

We compared allele distributions for autosomal variants ascertained for unrelated members of each collection (167 unrelated iPSCORE samples and 204 HipSci samples) after variant calling and filtering to ensure that differences between WGS from each collection did not create widespread systematic artifacts in variant calling. Allele distributions were compared between the studies using a chi-squared test with a Bonferroni correction. For instance, for an insertion, the number of samples with zero, one, or two copies of the insertion in iPSCORE were compared to the number of samples with zero, one, or two copies of the insertion in HipSci using the chi-squared test. Variants with Bonferroni-corrected p<0.05 were tagged in the VCF file. For this analysis, missing genotypes were also included as a unique allele when present.

We calculated Hardy Weinberg Equilibrium to identify variants that could be affected by batch effects in variant calling or that were poor quality. We used all unrelated blood/fibroblast samples and considered autosomal biallelic duplications and deletions from Genome STRiP (Handsaker et al., 2015) as well as all variant classes ascertained by SpeedSeq (Chiang et al., 2015) and MELT (Gardner et al., 2017). We tested HWE using a chi-squared test to compare the counts of the observed genotypes to those expected given HWE. SVs with Bonferroni corrected HWE p<0.05 were flagged as potentially not obeying HWE.

#### 3.1.3 Number of Calls ascertained Per Sample

Consistency in the number of non-reference calls per sample is associated with variant calls from high-quality WGS sequencing data, samples of similar ancestry, and algorithm performance. We counted the number of calls per sample for all algorithms to assess whether there were differences in the number of SVs identified in samples from each study, ancestry, or cell type from which the WGS was derived.

### 3.2 SpeedSeq

#### 3.2.1 Variant Calling

We used the split and discordant read pair-based structural variant caller LUMPY (v0.2.13)(Layer et al., 2014a) under its implementation in SpeedSeq (v0.1.2)(Chiang et al., 2015) to call duplications, deletions, inversions and other novel adjacencies referred to as “breakends” (BNDs). We ran LUMPY on each of 719 samples (478 from the HipSci collection and 478 from the iPSCORE collection) using the “speedseq sv” command with the -P option to retain probability curves in the output VCFs, -d to CNVnator (v0.3.3)(Abyzov et al., 2011) to calculate absolute copy number information on each sample, and -x to exclude a published list of genomic regions (ceph18.b37.lumpy.exclude.2014-01-15.bed) known to be potentially misassembled regions (Layer et al., 2014a; Li, 2014). Calls from individual samples were then genotyped using SVTyper (v0.1.4), before being combined into a single VCF file. Individual VCF files were sorted, and merged using svtools (v0.3.2) with the “sort” and “merge” command (slop 20bp) to remove overlapping breakpoints, resulting in a single VCF file with the most probable sites. Each sample was then genotyped at these merged sites using SVTyper and annotated with an absolute copy number using the “svtools copynumber” command. Variants were merged into a single VCF file, pruned, and reclassified under suggested parameters(Chiang et al., 2017). Individual VCFs were merged using “svtools vcfpaste” and further processed to remove additional identical variants using “svtools prune”. This set of breakpoints was then reclassified by using “svtools classify” to identify high confidence copy number variants by regressing the estimated copy number and “allele balance” information (non-reference/reference reads at an SV site) as well as to identify mobile element insertions in the reference genome (rMEI, which appear as deletions in our call set).

#### 3.2.2 Variant Processing

Because metrics such as replication rate may select variants that are reproducible artifacts, to remove as many known low-quality sites as possible, we first applied filtering guidelines suggested in a previous study (Chiang et al., 2017) as follows : 1) deletions that were less than 418bp were required to have split read support; 2) all non-BND variants were required to be at least 50bp in length; 3) BND calls required 25% percent support from either split or paired end reads; and 4) QUAL > 100 inversions were required to have at least 10% of evidence from split or paired end reads. Finally, to ensure a baseline level of genotyping consistency at each site, variants were filtered if they had a missing rate of > 10%.

#### 3.2.3 Variant Redundancy Collapsing

After running the Speedseq/SVtools pipelines and filtering variants as described above, the variant call set still contained overlapping variants suspected to be identical. To produce a single set of non-overlapping unique calls, we performed additional pruning steps. To identify and prune putatively identical calls that remained in our call set we implemented a graph-based approach: 1) we constructed a graph where SVs with reciprocal overlap of at least 50% are nodes connected by an edge; 2) we created a correlation matrix for each set of connected components using the allele balance (non-reference/reference reads at an SV site) at each site across individuals; 3) we refined the graph, retaining only the edges between SVs with r > 0.25 at a given site, which are likely to represent a single breakpoint; 4) we iterated through connected components, and chose variants with the highest median sample quality (MSQ) score, pruning other variants in the subgraph; and 5) in cases where one call was fully contained within another call and there was a correlation of at least 0.5 in allele balance between them, indicating that both calls were genotyped as non-reference in the same individual(s), only the site with the highest MSQ score was retained.

#### 3.2.4 Replication Rate Analysis and Filter Selection

During SpeedSeq quality analysis we investigated supporting reads (SU) and median sample quality (MSQ) as possible filtering criteria. MSQ was strongly associated with RR in iPSCORE twins and HipSci fibroblast iPSC pairs (Figure S2A, B) while the number of supporting reads was not (data not shown). For variant filtering, we determined variant class-specific MSQ thresholds, with the goal of ensuring at least 90% replication rate across all variant classes and retaining the maximal number of variants. Classes of variation that were highly reproducible before quality score filtering (>90% replication rate) were filtered at a 20 MSQ score, a threshold used in previous efforts (Chiang et al., 2017). With this approach, we performed additional filtering as follows: 1) deletions and rMEIs must have MSQ > 20; 2) duplications, inversions and BND calls must have MSQ > 100, 90, and 90 respectively. Deletions and rMEIs were genotyped most reproducibly, prior to filtering, while duplications were less reliably genotyped, reflecting the sensitivity of split read versus discordant read signal. After filtering, RR was on average 97% in twin pairs and slightly worse (92%) in fibroblasts-iPSC pairs (Figure S2C).

#### 3.2.5 Batch effect and Hardy Weinberg analysis

We tested variants on autosomes that passed the filters described above and 196 variants with missing rate > 10% but that otherwise passed filters for differences in allele distribution or deviations from HWE as described above (Section 3.1.2). We found that only 544 of 25,537 sites tested had different allele distributions (2.1%, Figure S2D). We also observed that 1,256 variants (4.9%) deviated from HWE, suggesting that batch effects do not affect SpeedSeq variant calls. We also observed that allele frequencies were highly correlated between variants detected in iPSCORE and HipSci.

#### 3.2.6 Calls Per Sample

After variant calling, we found that the number of SVs identified was consistent across samples, regardless of the study or cell type (Figure S2E-F). In agreement with previous SV discovery studies, we observed on average 10.2% more SpeedSeq variants per sample for those of African ancestry (4,260/sample) (Chiang et al., 2017; Sudmant et al., 2015) as compared to samples that were not predicted to be of African ancestry (3,863/sample).

### 3.3 Genome STRiP CNVDiscovery

#### 3.3.1 Variant Calling and Genotyping

Genome STRiP (svtoolkit 2.00.1611) CNVDiscovery (Handsaker et al., 2015), a population level read-depth based caller, was used to identify and genotype biallelic duplications and deletions as well as multiallelic CNVs (mCNVs) with suggested discovery parameters for deeply sequenced genomes (window size: 1000bp, window overlap: 500bp, minimum refined length: 500bp, boundary precision: 100bp, reference gap length: 1000). Because Genome STRiP is sensitive to differences in cell replication rate between samples derived from different cell types as well as in sequencing depth, we ran CNVDiscovery separately for iPSCORE fibroblast and blood samples and HipSci fibroblast samples. At the midstage of Genome STRiP discovery, 10 iPSCORE samples and 6 HipSci samples were removed from their respective discovery runs due to excessive variation in the number of calls per sample (exceeding the median call rate across all samples plus 3 median absolute deviations). To produce a call set where all sites were genotyped in all samples, sites discovered in either iPSCORE or HipSci samples were next genotyped using SVGenotyper in the opposite set (genotyping separately within these respective sets) and the combined list of discovered sites was genotyped in the remaining HipSci iPSC samples, which were excluded from discovery. Using this strategy, the Genome STRiP dataset was not biased by the presence of somatic CNVs in iPSCs, and differences due to WGS library preparation specific to each study were minimized. Additionally, output VCF files from genotyping each subset of samples were annotated to match those from variant discovery using the SVAnnotator (“-A CopyNumberClass, \ -A CNQuality \ -A VariantsPerSample \ -A NonVariant \ -A Redundancy”), to ensure that quality metric information was available for each variant within each subset of samples for downstream processing.

#### 3.3.2 Replication Rate analysis

A commonly suggested filtering parameter for SV detection is the per site quality score GSCNQUAL, described as being comparable for filtering of both duplication and deletion events (Andersson et al.). We thus tested the RR of Genome STRiP variants ascertained in iPSCORE samples as well as the replication of variants ascertained in the HipSci fibroblast samples (Figure S3A-B). Here we found that GSCNQUAL was highly correlated with RR in both twin pairs and iPSCs, but duplications and mCNVs had higher RR among twin pairs than iPSC-fibroblast pairs. Furthermore, deletions in both iPSCORE and HipSci sites were more reproducible under less stringent filtering than duplications and mCNVs. We selected 2, 12, and 14 as the minimum GSCNQUAL score required for deletions, mCNVs and duplications, respectively. We then filtered variants that were monoallelic in the data set as well as sites that had more than 10% of non-iPSC genotypes marked as low quality (LQ format field). These standard filters were applied before proceeding to combine the discovery sets of iPSCORE and HipSci and other data processing.

#### 3.3.3 Variant Redundancy Collapsing

To collapse redundant variants that were obtained through separate SV discovery for iPSCORE and HipSci samples, we first filtered the HipSci discovery set and the iPSCORE discovery set to those passing filters described above, and then intersected the call sets using bedtools(Quinlan, 2014; Quinlan and Hall, 2010). Overlapping sites were required to meet the following criteria in order to be considered redundant: 1) at least 50% reciprocal overlap; 2) Pearson correlation coefficient in the copy numbers of non-iPSC samples > 0.95; and 3) differences in less than 5% of non-mode genotypes in non-iPSC samples. To process these overlaps, we considered cases where two sites exactly overlapped (same coordinates), choosing the site with the largest sum of GSCNQUAL scores from iPSCORE and HipSci (Non-iPSC) samples sets as the high quality “primary” site and marking the other as “redundant”. Pairs of sites with exact overlaps were then removed from the analysis, and the remaining intersections were processed using a graph-based method similar to the one developed for Speedseq. Briefly, overlapping sites (nodes) were connected by edges weighted according to the average percentage overlap (the average of the percentage overlap of site B with A and the percentage overlap of site A with B) and which variant had the largest sum of GSCNQUAL scores from iPSCORE and HipSci (Non-iPSC) samples. Then, we iterated through connected components of the graph; chose the pair of sites that had the highest average overlap; and marked the variant with the largest sum of

GSCNQUAL scores as the “primary site” and the other variants in the cluster as redundant. For the X chromosome, the computation of correlation and differences among non-mode samples was done separately for males and females, requiring that sites pass criteria in males, females or both males and females, depending on whether each subgroup had variability. This was done to control for bias in correlation coefficients due to the difference in reference copy number for males and females on the X chromosome. Overall, this process resulted in 7,987 sites being reduced to 3,856 non-redundant primary sites.

#### 3.3.4 Stitching of CNVs

Genome STRiP occasionally reports a single CNV as several adjacent CNVs (Chiang et al., 2017). To address this issue, we analyzed sites that passed filtering, and were non-redundant, computing the correlation and distance between every pair of adjacent sites. We observed high genotype correlation between sites that overlapped or were close to each other (within ~40kb) (Figure S4A). Pairs of sites were considered for stitching into a single CNV if they had high overall correlation (*r* > 0.9) between copy number genotypes and at least 80% concordance between copy number genotypes of *non-mode* samples for each variant (union). Because variants that are very far from one another are less likely to be fragmented variant calls, we also selected a maximum distance between a pair of variants to consider for stitching. To do so, we examined the number and percentage of adjacent variant pairs that passed genotype correlation requirements at different distance thresholds, and selected 30 kb, which maximized the number and percentage of pairs passing these requirements (Figure S4B). We then identified correlated adjacent CNVs to be stitched using a graph-based method: 1) a genotype correlation matrix was created for all the CNVs on each chromosome using estimated copy numbers across samples; 2) a graph was drawn with CNVs as nodes, connecting a pair of CNVs with an edge if they resided on the same chromosome and had correlation from their copy number estimates >0.9; 3) for each connected component in the graph with more than a single CNV, CNVs were sorted by position and each adjacent pair was examined for potential stitching; and 4) CNVs were merged if they passed the correlation/concordance criteria described above and were within 30kb of one another. This approach ensured that only highly correlated adjacent CNVs were merged. In cases where a set of CNVs was chosen to be stitched, a new breakpoint spanning the start point of the first CNV to the end point of the last CNV (sorted by start point) was defined, referred to hereafter as the “stitch” breakpoint, while the other CNVs in the cluster were considered “constituent” sites. Note that in cases when a stitch cluster was made up of a single CNV containing one or more smaller CNVs, the large CNV was identified as a stitch breakpoint. Overall, this process lead to 3,558 sites being combined into 1,252 putative “stitch” breakpoints, 355 of which were large breakpoints in the call set that contained smaller breakpoints, and 897 were new breakpoints. The set of 897 new stitch breakpoints (not already genotyped in our set), were then genotyped across all samples using Genome STRiP SVGenotyper (CNVDiscovery), separately for iPSCORE samples, HipSci fibroblast samples, and HipSci iPSC samples (as was described in initial discovery 3.3.1). Finally, we compared the genotypes of the stitched breakpoint with the genotypes of the constituent sites, and those that did not have high correlation (average r < 0.9 across all constituents) were “unstitched”, and if the stitch breakpoint was one of the 897 new breakpoints genotyped, it was marked for filtering. If the new stitched breakpoint had over 10% low quality flagged genotypes (LQ) or was non polymorphic, the stitch cluster was also unstitched, and the breakpoint marked for filtering.

The vast majority of new stitch breakpoints were closely correlated with the constituents (862/897, 96%), suggesting that our stitching strategy indeed identified single CNVs that were broken into fragments (Figure S4C). An additional 7/862 correlated sites failed low quality genotype filtering criteria, yielding 855/897 (95%) new stitch breakpoints which passed all criteria. Overall, the process yielded 1207 unique sites (855 newly stitched sites and 353 sites that had been previously genotyped) comprised of 2-30 distinct CNVs each (Figure S4D). For analysis of the non-redundant set, we filtered these constituent sites and retained the stitch breakpoints. After the filtering, deduplication, and stitching process, remaining non-redundant variants had high replication fractions in each individual twin pair and fibroblast iSPC pair (Figure S4C) and high average replication rates on a per site basis (Figure 3A).

#### 3.3.5 Batch effect and Hardy Weinberg analysis

After filtering, variant collapsing and stitching, we tested for differences in allele distribution and deviations from HWE as described above (Section 3.1.2). Non-mode allele frequency was highly correlated between unrelated samples from iPSCORE and HipSci (Figure S4D) though a small number of variants (276/10,302 autosomal CNVs) were identified as having possible differences in allele distribution or deviation from HWE.

#### 3.3.6 Calls Per Sample

After variant calling and collapsing, we observed approximately the same number of calls per sample among iPSCORE and HipSci fibroblast samples, and no notable outliers among them (Figure S4E). As with other variant callers, we saw larger numbers of calls per sample among samples from the African predicted superpopulation (~28% more calls per sample).

Additionally, we found a small number of low quality genotypes per sample (Figure S4F) on the samples from which we performed discovery. HipSci iPSCs have higher rates of low quality genotypes because they were excluded from filtering that of sites based on their percentage of genotypes that were tagged as low quality (FORMAT = LQ) because they were genotyped separately and excluded from the CNVDiscovery pipeline. These results suggest that the discovery and genotyping approach was successful in preventing systematic batch effect variants.

### 3.4 Genome STRiP LCNVDiscovery

#### 3.4.1 Variant Calling

To identify CNVs longer than 100 kb, which we refer to as long CNVs (LCNVs) we used the LCNVDiscovery module of the Genome STRiP toolkit (svtoolkit 2.00.1611). This pipeline uses information from depth of coverage in fixed-size bins across the genome, and while sample normalization is performed across samples, individual samples are called separately. Prior to LCNVDiscovery, we generated depth profiles for all genomes using GenerateDepthProfiles with suggested parameters (maximumReferenceGapLength = 1000, profileBinSize = 10000). Then, similar to our approach in Genome STRiP CNVDiscovery, iPSCORE samples, Hipsci fibroblasts and HipSci iPSCs were processed separately when running the LCNVDiscovery module (maxDepth=50). We collected the calls from each sample and filtered them with the suggested parameters (NBINS ≥10 and a SCORE ≥1000). Sites that were entirely contained within the centromere or overlapped the entire centromere were removed and variant sites were required to have an absolute copy number greater than 2.75 or less than 1.25 for duplications and deletions, respectively (Figure S5A).

#### 3.4.2 Variant Processing and QC

Genome STRiP LCNVDiscovery identifies sites per individual sample, so it is necessary to identify redundant sites that are called in different samples. To find redundant CNVs representing a single breakpoint, sites with a reciprocal overlap of at least 80% were grouped into clusters and a single breakpoint spanning the minimum start position to the maximum end position of CNVs in the group was used to represent the merged site. Individual CNVs that were within these clusters were marked as merged constituents, and excluded from non-redundant set, while those that didn’t overlap with CNVs from another individual were considered unique variants that were present in only a single sample (Figure S5A-B). Absolute copy number estimates were rounded in order to produce integer copy number estimates similar to Genome STRiP CNVDiscovery. We identified 73 redundant sites comprised of 2 to 19 CNVs detected in individuals. On average, twin replication rates of the filtered variants was > 75% but very few large common variants were identified (Figure S5C). After filtering and collapsing variants, we obtained 432 unique LCNV sites, with 200 duplications, 166 deletions, and 66 mCNV (size range: 100 kb to 5 Mb, Figure S5D). On average each individual had 4 large duplications and 3 large deletions (Figure S5E).

### 3.5 MELT

#### 3.5.1 Variant Calling

Mobile element insertions (MEIs) were called using the Mobile Element Locator Tool (MELT) (Gardner et al., 2017). We used the MELT (v2.0.2) SPLIT workflow to discover, genotype, merge and annotate MEI calls for ALU, SVA and LINE1 elements. We also included discovered 1KGP MEI sites (Sudmant et al., 2015) as priors in “MELT GroupAnalysis”.

#### 3.5.2 Replication Rate Analysis and Filter Selection

While MELT does not output quantitative quality scores, it does flag variants that meet one or more of several criteria. These criteria include: 1) sites that overlap low complexity regions (lc), 2) have more than 25% missing genotypes (s25), 3) have a ratio of evidence for the left and right breakpoint (LP/RP) that is > 2 standard deviations from the ratio among all other sites (rSD), or 4) have a larger than expected number of discordant read pairs that are also split reads (hDP). We tested whether the flags, or combinations or flags, were associated with RR and found that filtering on all suggested criteria improved RR considerably for detected MEIs, raising it from below 0.6 to ~0.9 for ALU, LINE, and SVA elements (Figure S6A-B). Among these quality metrics, filtering on low complexity resulted in the best improvement compared with the other individual filters; however, filtering on all quality tags was necessary to improve RR to 0.9. Additionally, MELT outputs a quality tranche score from 1-5 (defined as “ASSESS”) that describes the types of evidence used to determine the location of the insertion site. For example, the highest quality insertion sites are given a score of 5, and has a target site duplication sequence flanking the MEI supported by split reads. Filtering with higher quality tranche score thresholds also improved replication rate, either before or after filtering using all flags (data not shown). We chose to filter variants that that were flagged for any criteria, and also required a quality tranche score of 5, for maximum stringency and best RR improvement. After filtering, individual twin and fibroblast-iPSC pairs had high replication percentages (>0.9 Figure S6C).

#### 3.5.3 Batch effect and Hardy Weinberg analysis

We tested all MELT variants for differences in allele distribution and deviation from HWE as described above (Section 3.1.2) and found that only 527/9,566 autosomal MEIs had differences in allele distribution (49/527) or showed deviation from HWE (492/527) (Figure S6D). Additionally, non-reference allele frequency in iPSCORE and HipSci was highly correlated (r > 0.9), suggesting batch effects did not influence MEI calls.

#### 3.5.4 Calls Per Sample

MELT variants were highly consistent in calls per sample in both studies, (mean 1,107 and 1,097 calls/sample in iPSCORE and HipSci fibroblast samples respectively) and in all cell types, while having very few missing genotypes (median 1/sample, Figure S6E-F). We observed an increased number of ALU, LINE1, and SVA elements per sample in samples from individuals of African ancestry (1,144 ALU/118 LINE1/53 SVA per sample versus 952 ALU/ 105 SVA/ 45 SVA sample for Non-African samples from iPSCORE).

### 3.6 HipSTR

#### 3.6.1 Variant Calling

Short tandem repeat (STR) variants were genotyped using the HipSTR algorithm (v0.5.61) (Willems et al., 2017), on a set of 1,527,077 GRCh37 autosomal STR regions that were provided by the tool (https://github.com/HipSTR-Tool/HipSTR-references/raw/master/human/GRCh37.hipstr_reference.bed.gz). Because only HipSci WGS data was PCR-free, special considerations were required to run HipSTR, as it uses PCR stuttering models to genotype repeats and assumes all WGS samples were generated using the same pipeline. For STR genotyping, PCR-free data produces more accurate genotypes, thus we first ran HipSTR at STR sites in all 446 HipSci samples under standard settings. Next, we genotyped the iPSCORE samples using the HipSci genotypes as references (--ref option). Finally, we genotyped iPSCORE samples separately without using the HipSci genotypes as reference alleles. We used only the diploid genotype option, as we lacked phased SNVs for all samples.

#### 3.6.2 Filtering and Preliminary Replication Rate Analysis

To filter HipSTR variants, we first used the supplied “filter_vcf.py” script with recommended thresholds for individual genotypes (min-call-qual = 0.9, max-call-flank-indel = 0.15, max-call-stutter = 0.15, --min-call-allele-bias= -2, min-call-strand-bias= -2). This procedure converts genotypes that do not pass these thresholds to “missing”. We examined the number of variant calls per sample and the number of missing genotypes when variants were genotyped in iPSCORE, iPSCORE using HipSci reference alleles, and in HipSci samples (Figure S16). Among iPSCORE samples, we observed a median of 122,249 calls per sample in African ancestry individuals and 111,613 calls per samples in non-African ancestry individuals (Figure S16A-D). While four samples from non-African ancestry individuals had a surprisingly large number of STRs, all but one individual self-reported as having partial African ancestry (Figure S16D). iPSCORE genotypes at HipSci reference alleles had similar numbers of calls per sample (median 110,023/sample) compared to the genotypes from iPSCORE alone (Figure S16E). African ancestry samples, however, had a smaller number of calls using the HipSci reference alleles likely because HipSci did not include African ancestry samples, so the African samples in iPSCORE were only genotyped for STRs discovered in Europeans. HipSci samples had about twice as many calls per sample (222,321/sample for HipSci fibroblast samples) compared to iPSCORE and fewer missing calls per sample, demonstrating that using PCR-free WGS provides better accuracy for STR genotyping. To obtain a high-quality set of STRs, we required >80% call rate for variants from each subset. We excluded one iPSCORE sample from this missingness calculation that had more than 70,000 missing calls. This filter resulted in high replication rates (> 92%) in each twin pair for both genotyping methods in iPSCORE, and even higher replication rates (>95%) in fibroblast-iPSC pairs for HipSci genotyping likely due to more accurate STR genotyping in the PCR-free WGS (Figure S17). Overall, the replication rate before all filtering and after processing improved from ~78% to ~94.4% in iPSCORE twins (Figure 3B).

#### 3.6.3 Combining the iPSCORE and HipSci data

HipSTR genotypes were combined between iPSCORE and HipSci by creating a single combined VCF file using the HipSci genotypes and iPSCORE genotypes at HipSci alleles. We additionally added iPSCORE genotypes for STRs that were unique to iPSCORE to the VCF file.

### 3.7 Unifying SpeedSeq and Genome STRiP CNVDiscovery and LCNVDiscovery Call Sets

Since different variant callers may detect the same variants using different methods, we developed a strategy to integrate variants from Genome STRiP and SpeedSeq call sets that were likely to represent the same site. To approach this problem, we used a graph-based method similar to those used to identify duplicates within SpeedSeq and Genome STRiP prior to this step (Sections 3.2.3 and 3.3.3). To generate clusters of overlapping SVs, we first intersected our filtered Genome STRiP calls (redundant sites removed, GSCNQUAL filtered, stitching sites included, stitched constituents excluded) with filtered SpeedSeq variants (redundant sites removed, standard filters, MSQ filtered) and retained all SV pairs with >50% reciprocal overlap or where one site completely contains another. SV pairs were required to have the same SV types, with exception being that mCNVs were allowed to match with both duplications and deletions and deletions were allowed to match with rMEI (as they appear as deletions). We built a graph where edges were represented by connected SV pairs that pass these overlap thresholds and SV type compatibility parameters. We iterated through connected components, testing every combination of elements in each connected component, and generating a new graph, connecting pairs of variants if they passed correlation thresholds between copy number genotypes (Genome STRiP) variants or allele balance ratios (SpeedSeq) at the sites. If the connected component contained a duplication and deletion from SpeedSeq and an mCNV from Genome STRiP, SV pairs were allowed to connect if their genotype evidence had a correlation 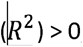, while other components required an 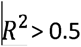. We then iterated through connected components of this new graph and selected the highest degree variant (connected to the most other variants) from each caller with the highest quality score (GSCNQUAL for Genome STRiP, MSQ for SpeedSeq) from which we chose one variant as the “primary” variant and all other variants as “secondary.” All variants in each cluster were marked with a cluster ID. In cases where a Genome STRiP deletion overlapped a SpeedSeq rMEI, the SpeedSeq variant was chosen as the primary site, and the Genome STRiP variant was assigned as secondary. In all other scenarios, the Genome STRiP variant was chosen as the primary variant and the SpeedSeq variant was the secondary due to the comparably higher replication rates for Genome STRiP variants and the granularity of having integer copy numbers.

This method assures that highly correlated variants with significant overlap are clustered together, and that generally, the larger, higher quality variants are chosen as representative primary sites. Sites that were assigned as primary sites from the intersection clusters, as well as unique variants from either variant call set that were not included in intersection clusters, were then selected to produce a non-redundant set of sites necessary for global analyses of SVs (Figure 4,5).

### 3.8 Comparison to SV Genotypes from Arrays

To estimate the false discovery rate (FDR) of the merged CNV call set we used 216 MEGA_Consortium_v2 arrays available for iPSCORE samples to perform an intensity rank sum (IRS) test to assess whether the SV genotypes after filtering agree with genotypes from array data. SNP arrays were analyzed using the Illumina GenomeStudio software (v2011.1) and were required to have an overall call rate of <97%. The log(R ratio) was obtained from the final report. We used the Genome STRiP Intensity Rank Sum Annotator to compare genotypes for a subset of the SV calls that were present in the 216 samples for which we had array data using the log R ratio as input. Before testing, the intensity matrix was first adjusted for covariates by regressing out the effects of batch and plate on a probe-wise basis using the statsmodels (v0.9.0) linear regression module. To assess our filtering strategy we tested 2,563 /15,437 SpeedSeq duplications and deletions, and 4,233/18,171 Genome STRiP CNVs that were present in at least one of the 216 individuals (before any filtering) and contained at least 3 probes and computed IRS FDR as in 1KGP (Sudmant et al., 2015). Restricting our analysis to 2,376 filtered and deduplicated SpeedSeq variants with array probes, we observed that deletions and duplications had an FDR of 5.35% and 3% respectively. Similarly, among 1,848 filtered and deduplicated Genome STRiP variants containing array probes, we observed that deletions, duplications and mCNVs had an FDR of 5.4%, 7.8%, and 7% respectively. These FDR estimates were similar to those in 1KGP and GTEx, although the probe density of arrays limited the number of sites we could test.

### 3.9 Comparison of i2QTL SVs to 1000 Genomes Project and GTEx SV Call-Sets

To investigate the quality and completeness of our SV calls, we compared them to GTEx v6p SV calls (Chiang et al., 2017) which used 147 deeply sequenced whole genomes (median 49.9X depth), and the robustly characterized 1000 Genomes Project Phase-3 call-set (Sudmant et al., 2015) derived from 2,504 shallowly sequenced samples (7.4X depth). While the GTEx call-set contains relatively few samples, the whole genome sequencing data and variant calling approach were similar to the approach used in i2QTL (Genome STRiP and SpeedSeq), and were thus used as a benchmark. Before analysis, we obtained VCF files with genotypes from 1KGP phase 3 (link?) and GTEx V6p (dbGaP accession number phs000424.v7.p1). Phased genotypes from 1KGP SVs were converted to unphased genotypes using the alternative allele information to enable comparison with the unphased SVs from i2QTL and GTEx. This enabled us to compute non-mode allele frequency for 1KGP and GTEx SVs to match the frequency measures used in this study. Because of the significant diversity of the 1KGP cohort (26 populations, 70% European) as compared to i2QTL (6 subpopulations, 80% European), we filtered the 1KGP data to 1,755 European samples, and used variants present in at least one of these samples. For co-discovery analyses, we used non-redundant sites from i2QTL as well as variants that passed filters and were part of redundancy clusters to maximize the potential overlap between sets. To identify putative co-discovered sites between i2QTL and either GTEx or 1KGP, CNVs (DUP, DEL, mCNV), rMEI and inversions from each call-set were intersected using “bedtools intersect” and co-discovered sites were selected using the following approach: 1) excluding inversions, all variants were required to have at least 25% reciprocal overlap, or if one variant was fully contained within the other, it was required to span at least 20% of the larger variant; inversions were required to have 80% reciprocal overlap; 2) variant classes were required to match with the exception of mCNVs, which were allowed to match with either duplications or deletions; for BND sites, we considered breakpoints within 50bp of each other to be matching; and 3) because we included 1KGP MEIs as priors in our MELT pipeline, MEIs co-discovered with 1KGP were known, and did not require overlap analysis. For overlap reported with i2QTL, we computed the fraction of sites co-discovered by one or both call-sets, considering non-redundant clusters a single site.

## 4 LD Tagging

For each of the 42,921 total non-redundant SVs and STRs that were within 1MB of an expressed gene in iPSCs (Jakubosky et al., 2019), we used bcftools (Li et al., 2009) to extract all SNPs 50 kb upstream and downstream. For each SV or STR, we calculated LD as the correlation (Pearson R^2^) with the genotypes of each surrounding SNV or indel genotyped in i2QTL WGS and selected the variant with the strongest LD.

## Acknowledgments

This work was supported in part by the National Science Foundation, CIRM grant GC1R-06673-B, and NIH grants HG008118, HL107442, DK105541 and DK112155. D.A.J. was supported by the National Library Of Medicine of the National Institutes of Health under Award Number T15LM011271. W.W.YG. was supported by the National Heart, Lung, And Blood Institute of the National Institutes of Health under Award Number F31HL142151. M.J.B. was supported by a fellowship from the EMBL Interdisciplinary Postdoc (EI3POD) program under Marie Skłodowska-Curie Actions COFUND (grant number 664726). S.B.M. was supported by NIH grant U01HG009431.

## Author Contributions

Conceptualization, D.A.J, E.N.S, M.D., O.S., S.B.M., C.D and K.A.F.; Methodology, D.A.J, E.N.S., M.D., M.J.B., W.W.Y.G., O.S., S.B.M; Formal Analysis, D.A.J.; Data Curation, D.A.J., A.C.D., H.M.; Writing – Original Draft, D.A.J, E.N.S, M.D., C.D. and K.A.F.; Visualization, D.A.J; Supervision, O.S., S.B.M and K.A.F.; Funding Acquisition K.A.F., O.S. and S.B.M.

## Supplemental Tables

**Table S1: Subject Information.** Phenotypic information about 477 individuals from iPSCORE and HipSci with WGS used in this study including age, sex, predicted superpopulation, reported and annotation of family, twin status, presence in the unrelated set.

**Table S2: WGS information.** Information describing WGS samples included in variant calling including cell type, subject, study, and also median coverage for each genome.

**Table S3.** Filtering parameters overview. Table describing the filtering parameters used when generating the i2QTL SV/STR call set, stratified by variant caller and variant class when necessary. The “Applied” column indicates whether the filter was applied to set specific low-quality genotypes to missing on a per sample basis within each variant class (“Per Genotype” filters) or whether the filter was used to filter an entire site for removal from downstream analysis (“Per Site” filtering). The column “Filtering Parameter” indicates the specific filter,

| Variant Caller | Variant Class | Applied | Filtering Parameter | Threshold | Rationale |
| --- | --- | --- | --- | --- | --- |
| Genome STRiP<br>CNVDiscovery | DEL | Per Site | GSCNQUAL | 2 | RR |
|  | DUP | Per Site | GSCNQUAL | 14 | RR |
|  | mCNV | Per Site | GSCNQUAL | 12 | RR |
|  | ALL | Per Site | % Low Quality Genotypes (LQ tag) | < 10% |  |
| Genome STRiP<br>LCNVDiscovery | ALL | Per Site | Contained in Centromere | N/A | Suggested (Methods) |
| | ALL | Per Site | NBINS | $\geq 10$ | Suggested (Methods) |
| | ALL | Per Site | SCORE | $\geq 1000$ | Suggested (Methods) |
| | ALL | Per Genotype | Absolute Copy Number | $1.25 < \text{or} > 2.75$ | Suggested (Methods) |
| SpeedSeq | DEL | Per Site | MSQ | 20 | RR |
|  | DUP | Per Site | MSQ | 100 | RR |
|  | rMEI | Per Site | MSQ | 20 | RR |
|  | INV | Per Site | MSQ | 90 | RR |
|  | BND | Per Site | MSQ | 90 | RR |
| | BND | Per Site | $\geq 25\%$ PE/SR support | N/A | Chiang et al. 2017 |
|  | INV | Per Site | QUAL | >100 | Chiang et al. 2017 |
| | INV | Per Site | $\geq 10\%$ PE or SR support | N/A | Chiang et al. 2017 |
|  | ALL | Per Site | % Missing | < 10% |  |
| MELT | ALL | Per Site | low complexity (lc) | N/A | RR/Gardner et al. 2017 |
|  | ALL | Per Site | >25% missing (s25) | N/A | RR/Gardner et al. 2017 |
|  | ALL | Per Site | LP/ RP > 2.0 standard deviations (rSD) | N/A | RR/Gardner et al. 2017 |
|  | ALL | Per Site | more discordant pairs are also split than expected (hDP) | N/A | RR/Gardner et al. 2017 |
|  | ALL | Per Site | ASSESS | =5 | RR |
| HipSTR | ALL | Per Genotype | call quality (Q) | 0.9 | RR/Willems et al. 2017 |
| | ALL | Per Genotype | Fraction reads with flanking indel (DFLANKINDEL/DP) | $\leq 0.15$ | |
| | ALL | Per Genotype | Fraction Reads With Stutter (DSTUTTER/DP) | $\leq 0.15$ | |
|  | ALL | Per Genotype | log10 allele bias p-value (AB) | >-2 |  |
|  | ALL | Per Genotype | log10 strand bias p-value (FS) | >-2 |  |

**Table S4 Non redundant structural variants.** Table describing the non-redundant variants including position, variant class, variant caller, and evidence supporting the site.

## Supplemental Figures

**Figure S1.**
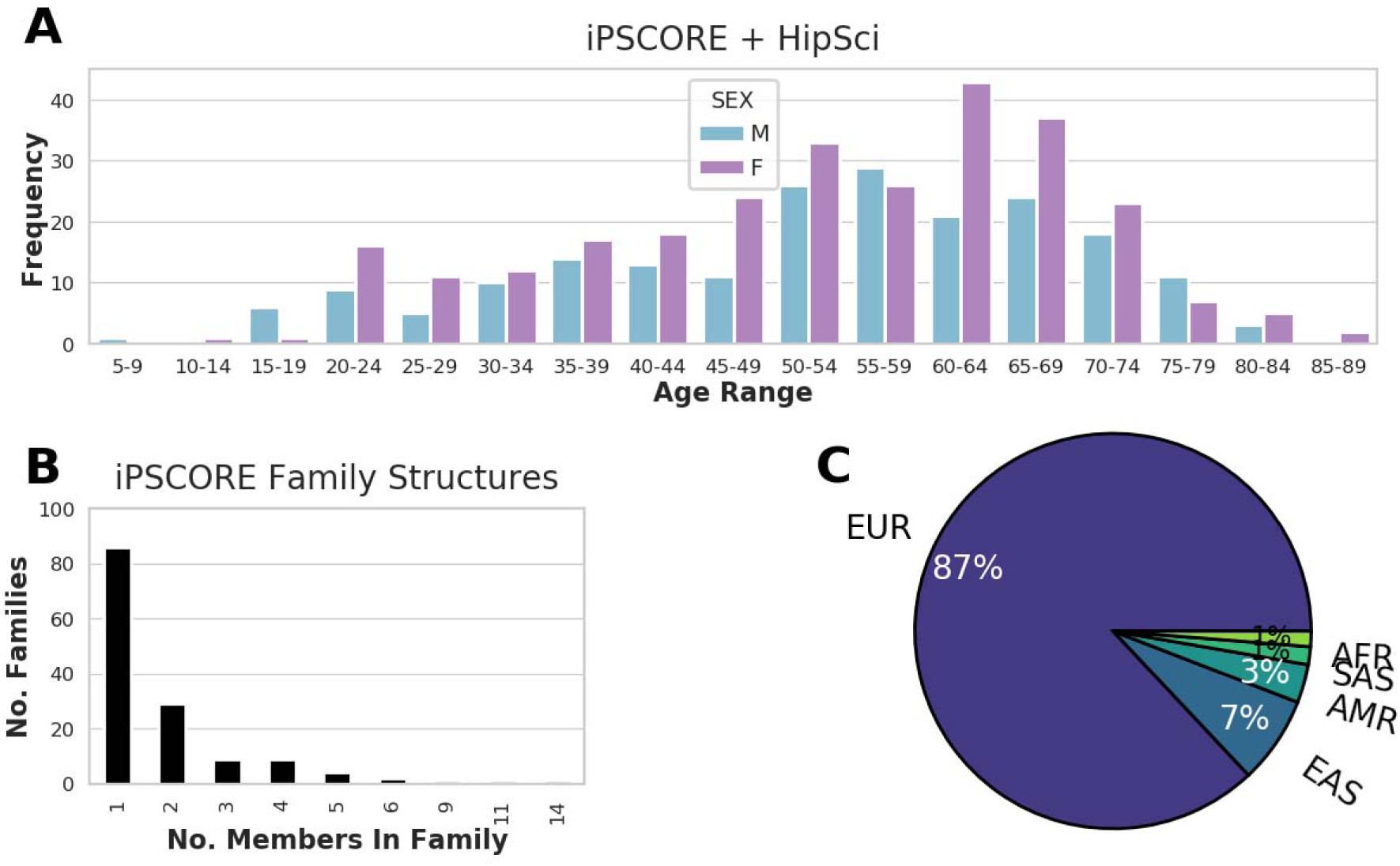
i2QTL Subject Information. (A) Age distribution of WGS donors stratified by sex. (B) Size of families in iPSCORE. (C) Number of individuals from iPSCORE and HipSci assigned to each of the 1000 Genomes Project superpopulations using genotype data (Auton et al., 2015).

**Figure S2:**
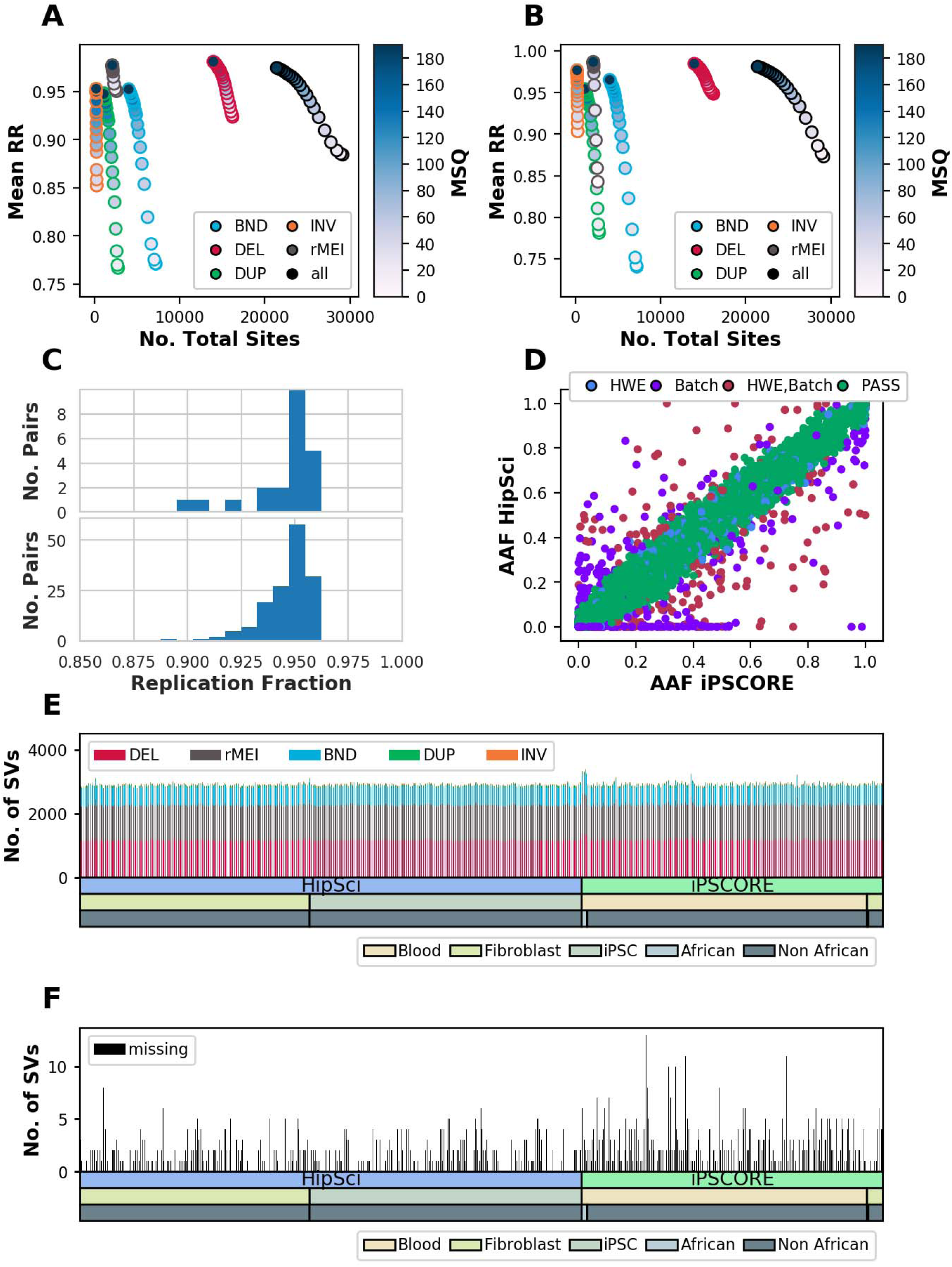
SpeedSeq Quality Control. (A and B) Replication rate in (A) iPSCORE monozygotic twins and (B) HipSci fibroblast iPSC pairs as a function of the number of total sites that pass filtering thresholds for median sample quality score (MSQ). (C) Replication rate distribution in monozygotic twin pairs (upper) and fibroblast iPSC pairs (lower). (D) Comparison of non-reference allele frequency of calls in iPSCORE unrelated samples and HipSci fibroblast samples, colored by whether their genotype distributions were flagged for deviation from Hardy Weinberg Equilibrium (blue) or potential systematic differences between genotypes in HipSci and genotypes in iPSCORE (“Batch”, purple), or both (red). (E) Number of events per individual after filtering for each variant type. (F) Number of events with a missing genotype after filtering.

**Figure S3:**
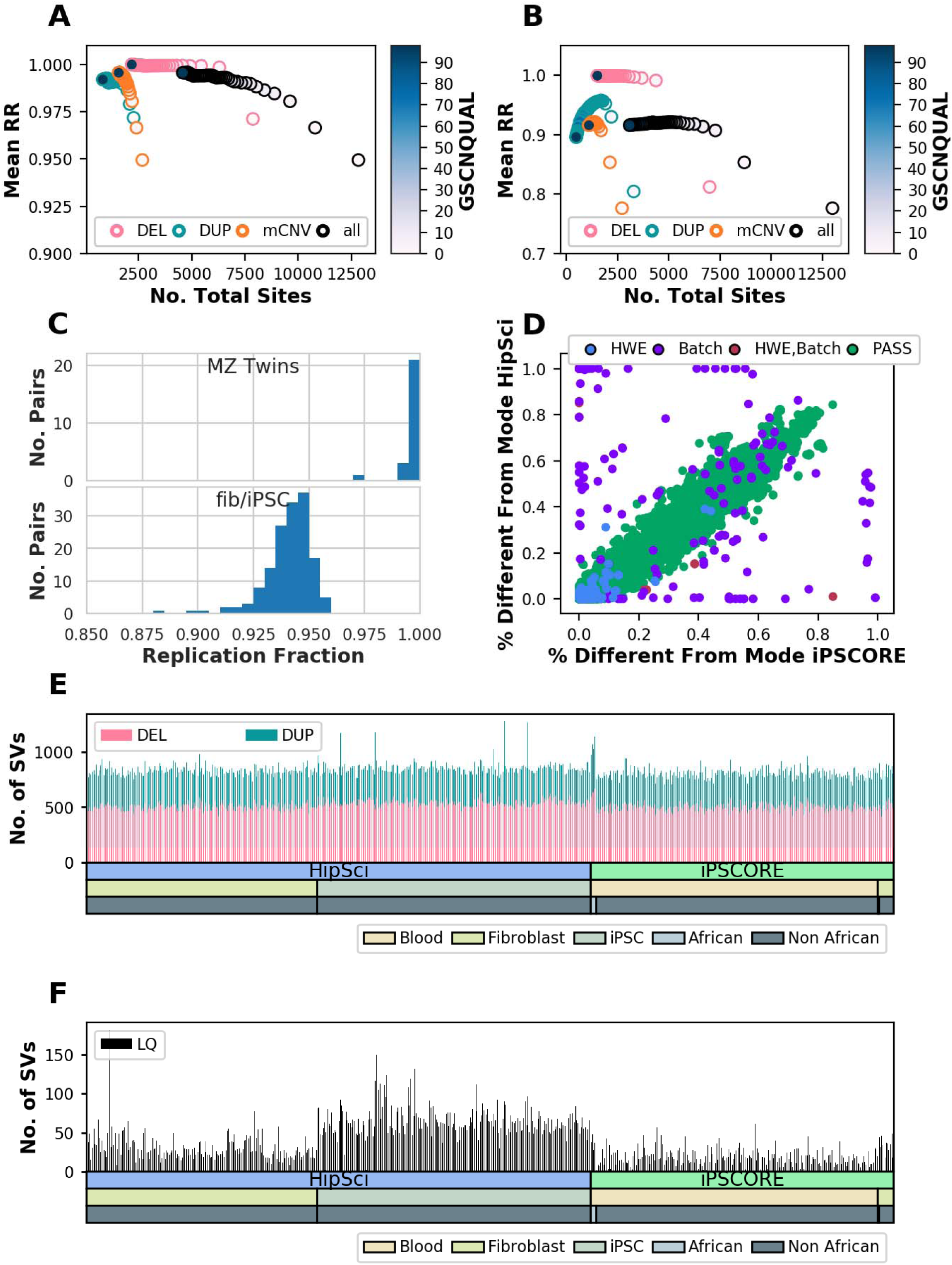
Genome STRiP Quality Control. (A, B) Replication rate in (A) iPSCORE monozygotic twins and (B) HipSci fibroblast iPSC pairs as a function of the number of total sites that pass filtering thresholds for GSCNQUAL. (C) Replication rate distribution in monozygotic twin pairs (upper) and fibroblast iPSC pairs (lower). (D) Comparison of percent of samples different from the copy-number mode in iPSCORE unrelated samples and HipSci fibroblast samples, colored by whether their genotype distributions were flagged for deviation from Hardy Weinberg Equilibrium (blue) or potential systematic differences between genotypes in HipSci and genotypes in iPSCORE (“Batch”, purple), or both (red). (E) Number of duplication or deletion events per individual after filtering. (F) Number of events with a genotype tagged as LQ (low quality) after filtering. Note that we filtered to variants that were less than 10% LQ rate among iPSCORE samples and HipSci fibroblast samples. Therefore, we observed more LQ sites among HipSci iPSCs.

**Figure S4:**
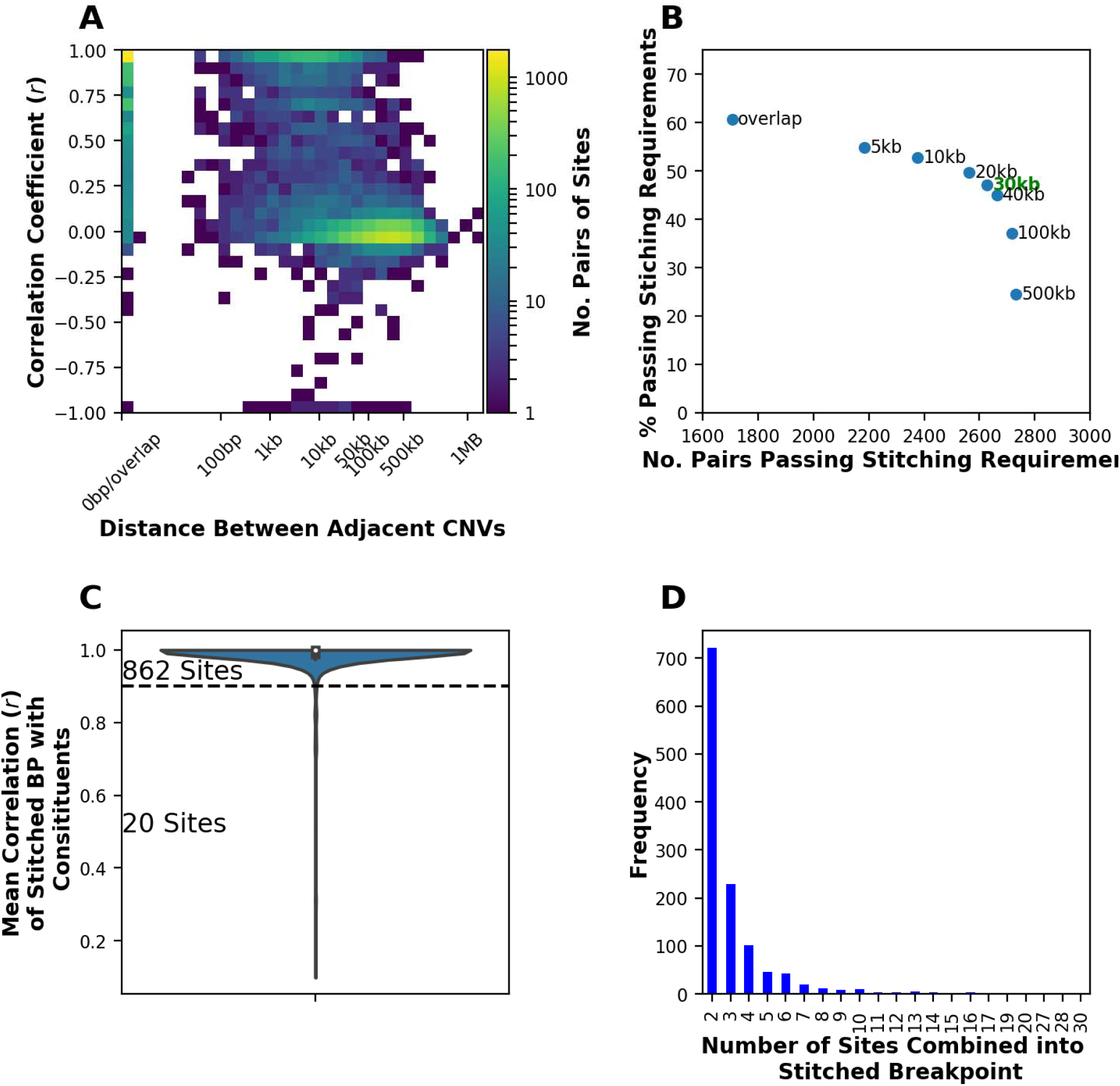
Genome STRiP Stitching. (A) 2D histogram showing the distance and correlation between pairs of adjacent CNVs, colored by the number of pairs of sites. This information was used to decide the maximum distance threshold used to stitch neighboring sites. (B) Number of pairs that passed stitching requirements versus the percentage which passed stitching requirements for different thresholds of maximum distance between variants (see Methods). We chose a threshold of 30kb (green) as this maximized the percent of pairs that passed stitching requirements, and few pairs of sites greater than 30kb apart were correlated. (C) Mean correlation of the stitched breakpoint genotypes with the genotypes of the constituent sites. In cases where the stitched breakpoint correlated at less than 0.9, it was discarded and the site was “unstitched”. Stitch breakpoints were also unstitched if more than 10% of samples had low quality (LQ) genotypes among the 477 iPSCORE and HipSci (fibroblast) samples (D) Number of sites combined into stitched breakpoints.

**Figure S5.**
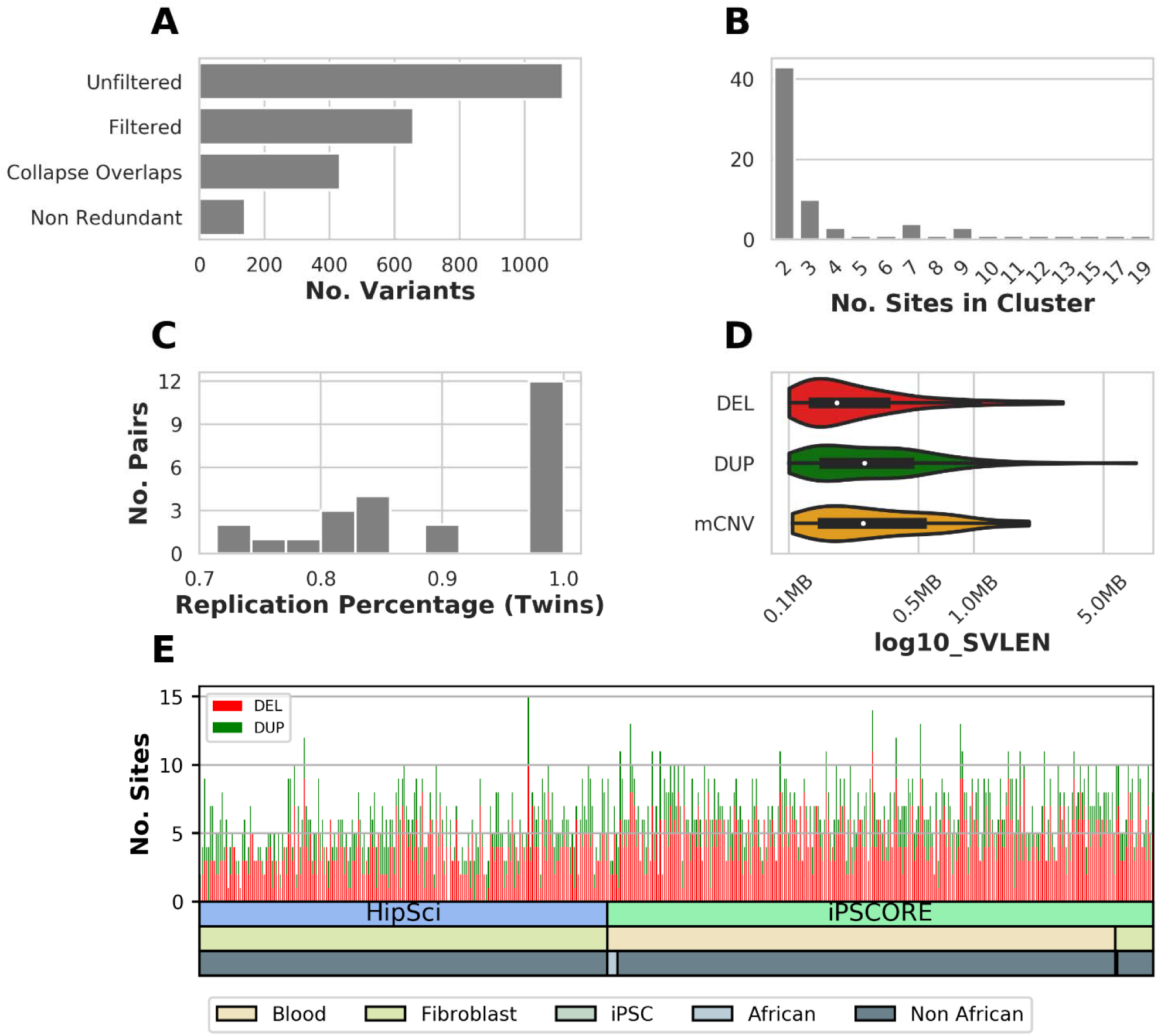
Genome STRiP LCNV Quality Control and Filtering. (A) Number of variants identified by the Genome STRiP LCNV pipeline at each filtering step. (B) Because GS LCNV variants are detected separately for each sample, all breakpoints are not genotyped in each sample, and large common variants may have different coordinates in distinct samples. Therefore, we collapsed all variants with reciprocal overlap >80% into single sites. Here we show the number of clusters of overlapping variants and the number of sites per cluster. (C) Replication percentage in each iPSCORE twin pair (D) Length distribution and (E) number of duplications and deletions per sample for variants after filtering and collapsing.

**Figure S6:**
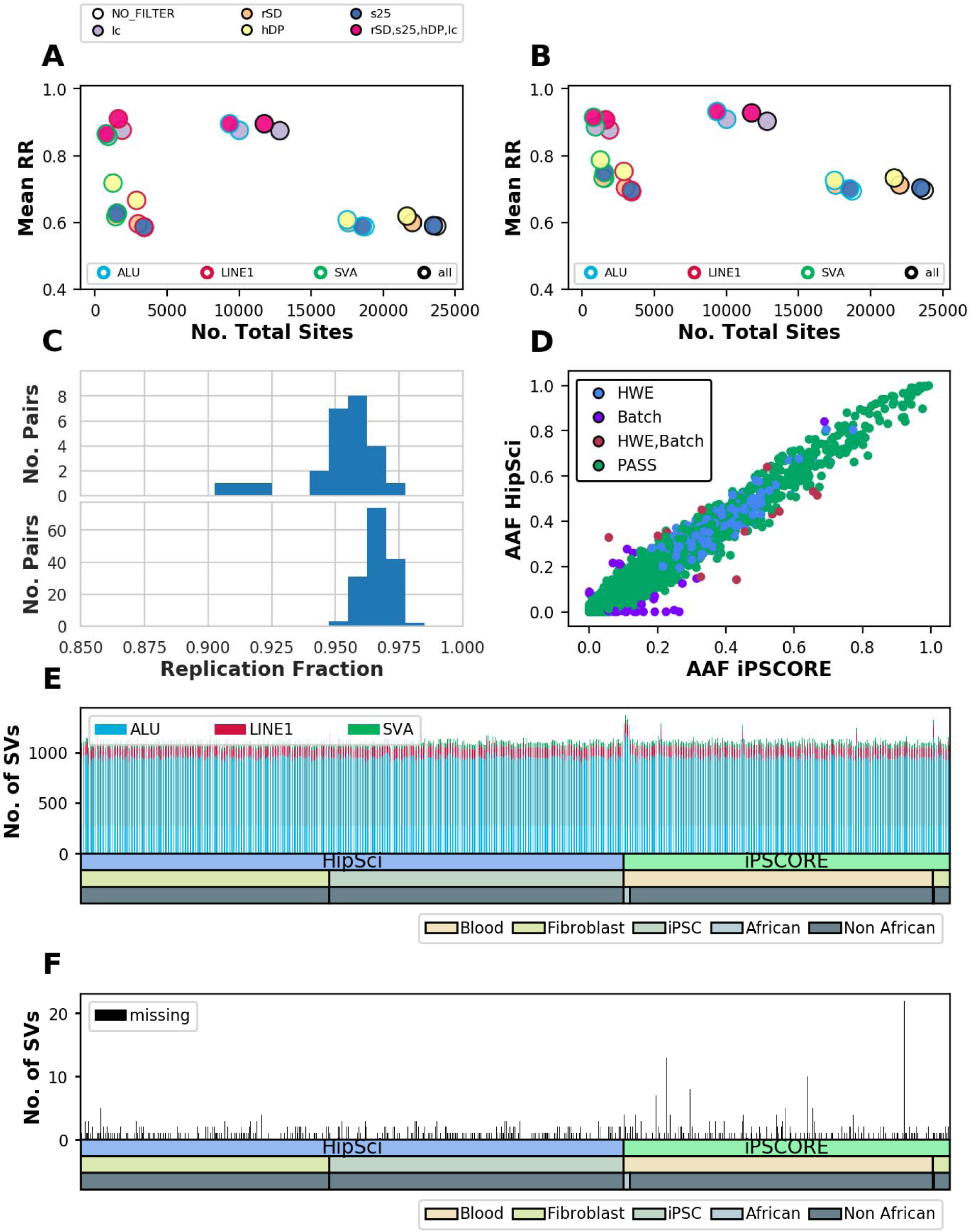
MELT Quality Control. (A and B) Replication rate in (A) iPSCORE monozygotic twins and (B) HipSci fibroblast iPSC pairs as a function of the number of total sites that pass filtering thresholds various parameters that are suggested by MELT, colors show how many of these filters were applied. (C) Replication rate distribution in monozygotic twin pairs (upper) and fibroblast iPSC pairs (lower). (D) Comparison of non-reference allele frequency of calls in iPSCORE unrelated samples and HipSci fibroblast samples, colored by whether their genotype distributions were flagged for deviation from Hardy Weinberg Equilibrium (blue) or potential systematic differences between genotypes in HipSci and genotypes in iPSCORE (“Batch”, purple), or both (red). (E) Number of events per individual after filtering for each variant type. (F) Number of events with a missing genotype after filtering.

**Figure S7.**
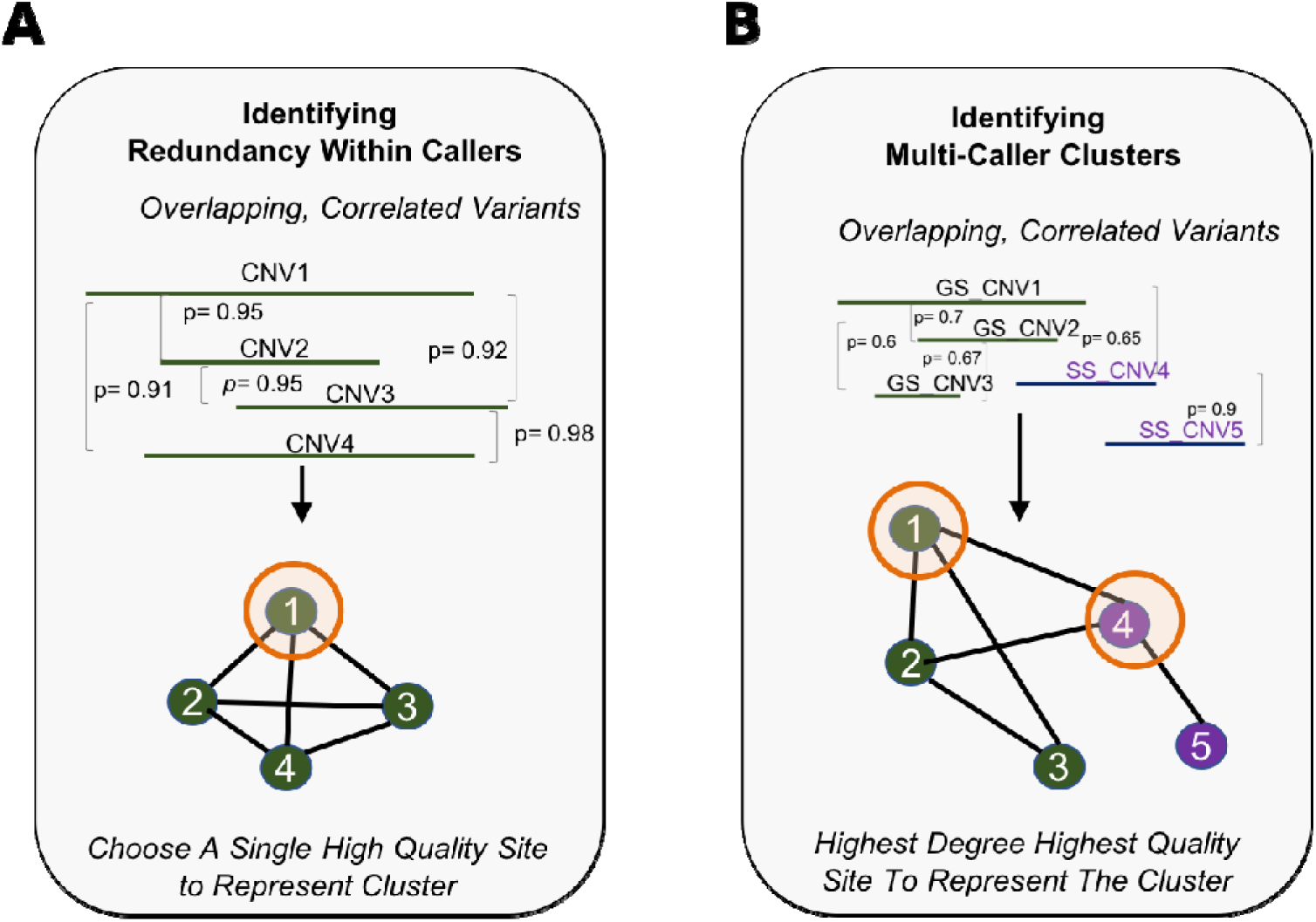
Collapsing Redundancies Within and Between Variant Callers. (A) Strategy for collapsing redundancies between callers by identifying clusters of overlapping variants with high correlation between variants. While the approach for SpeedSeq and Genome STRiP was slightly different (Methods), overall, the goal was to select a single high-quality site to represent a cluster of correlated variants. (B) Illustration of the strategy for identifying redundant sites called by multiple algorithms. Here, similar to collapsing variants within caller, overlapping variants from different algorithms that had correlated genotyping information were represented as a graph. Edges were drawn between variants that did not overlap if genotype correlation was above a specific threshold (Methods), and the highest degree, highest quality variant was chosen from either Genome STRiP or SpeedSeq to represent the cluster.

**Figure S8.**
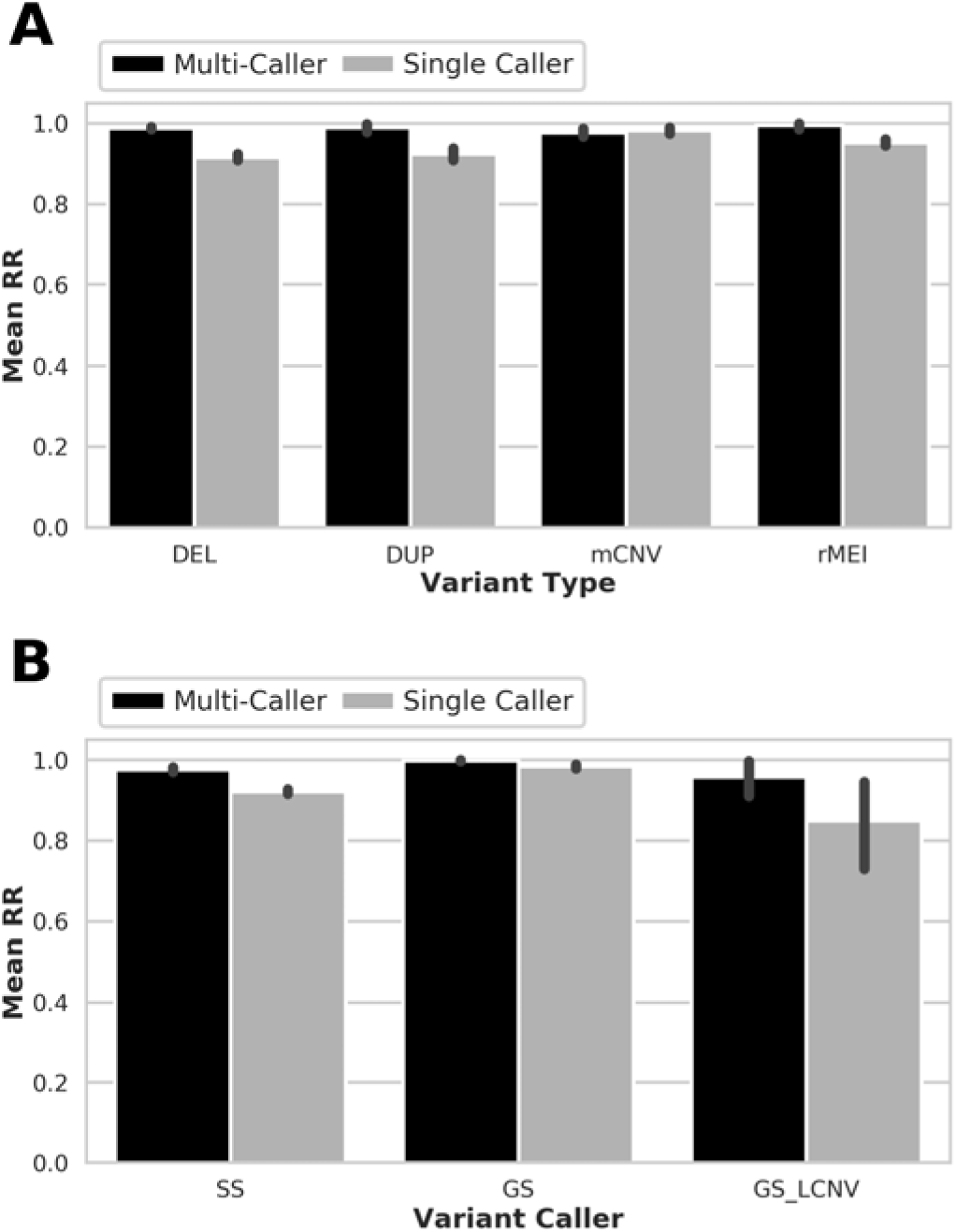
Replication rate of variants that are identified by multiple callers. (A, B) Average replication rate of variants that were or were not discovered by multiple callers stratified by (A) variant type or (B) variant caller.

**Figure S9.**
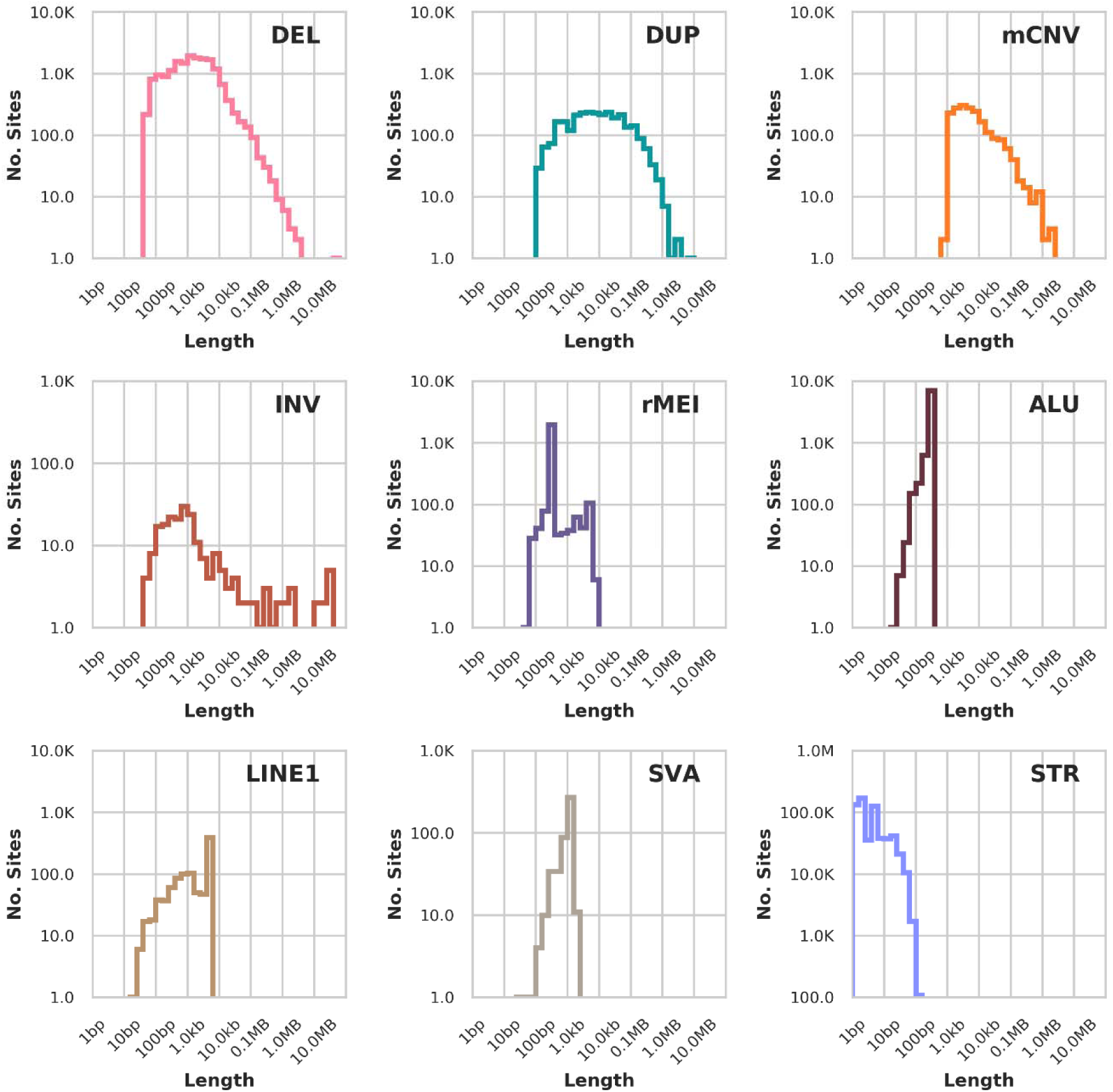
Length Distribution of Non-Redundant Variants. Length distribution of variants in the non-redundant call set as identified in 477 individuals. The length for STRs is calculated as the maximum absolute difference in base pairs from the reference allele at a site while the length for CNVs (DEL, DUP, mCNV) is the size of a single copy unit. For rMEI, ALU, LINE1, and SVA the length represents the estimated insertion size of the variant. For inversions, the length is the distance between the two breakpoints.

**Figure 10.**
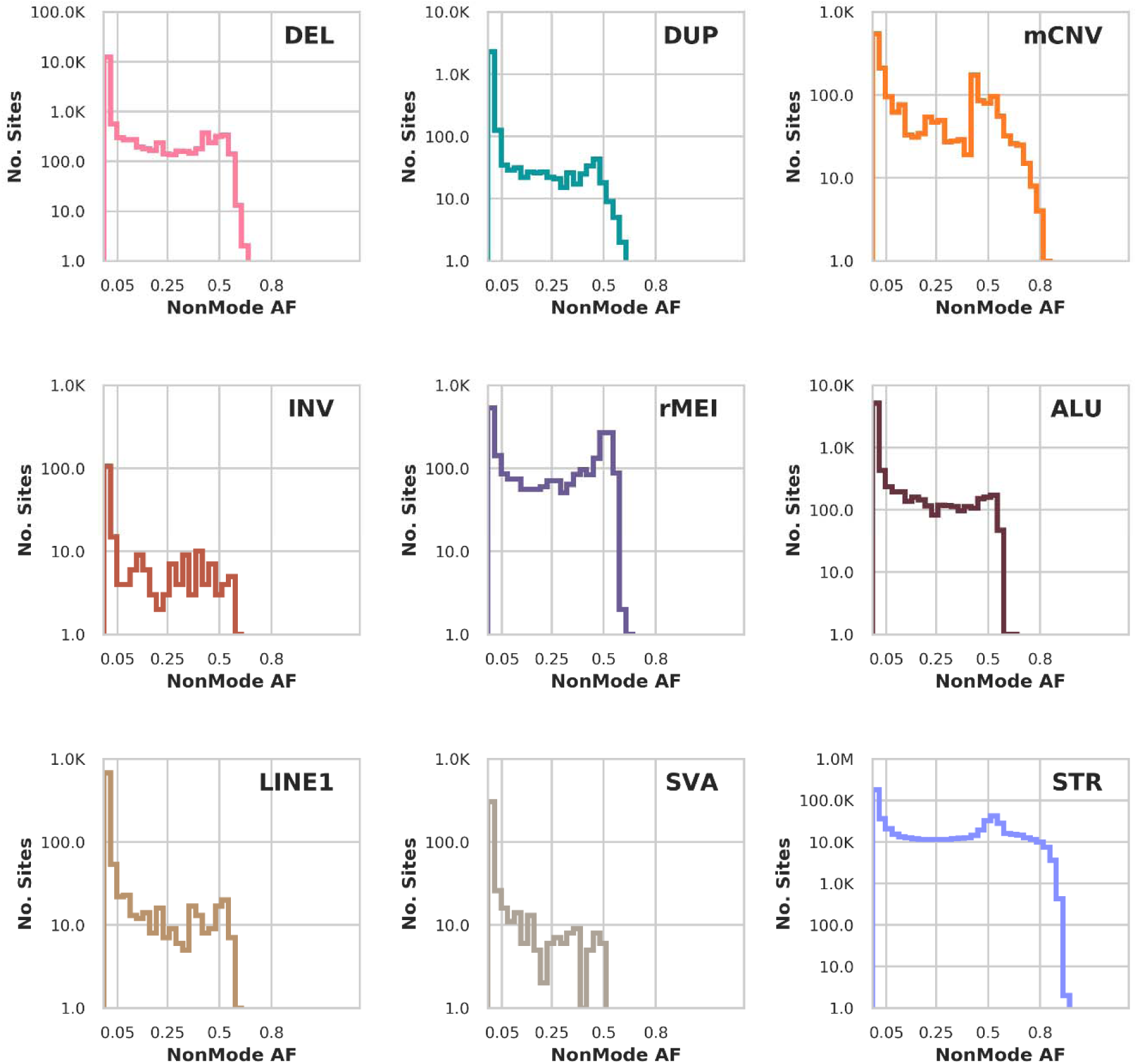
Allele Frequency Distribution of Non-Redundant Variants. Distribution of the non-mode allele frequency in each variant class for i2QTL unrelated samples.

**Figure S11.**
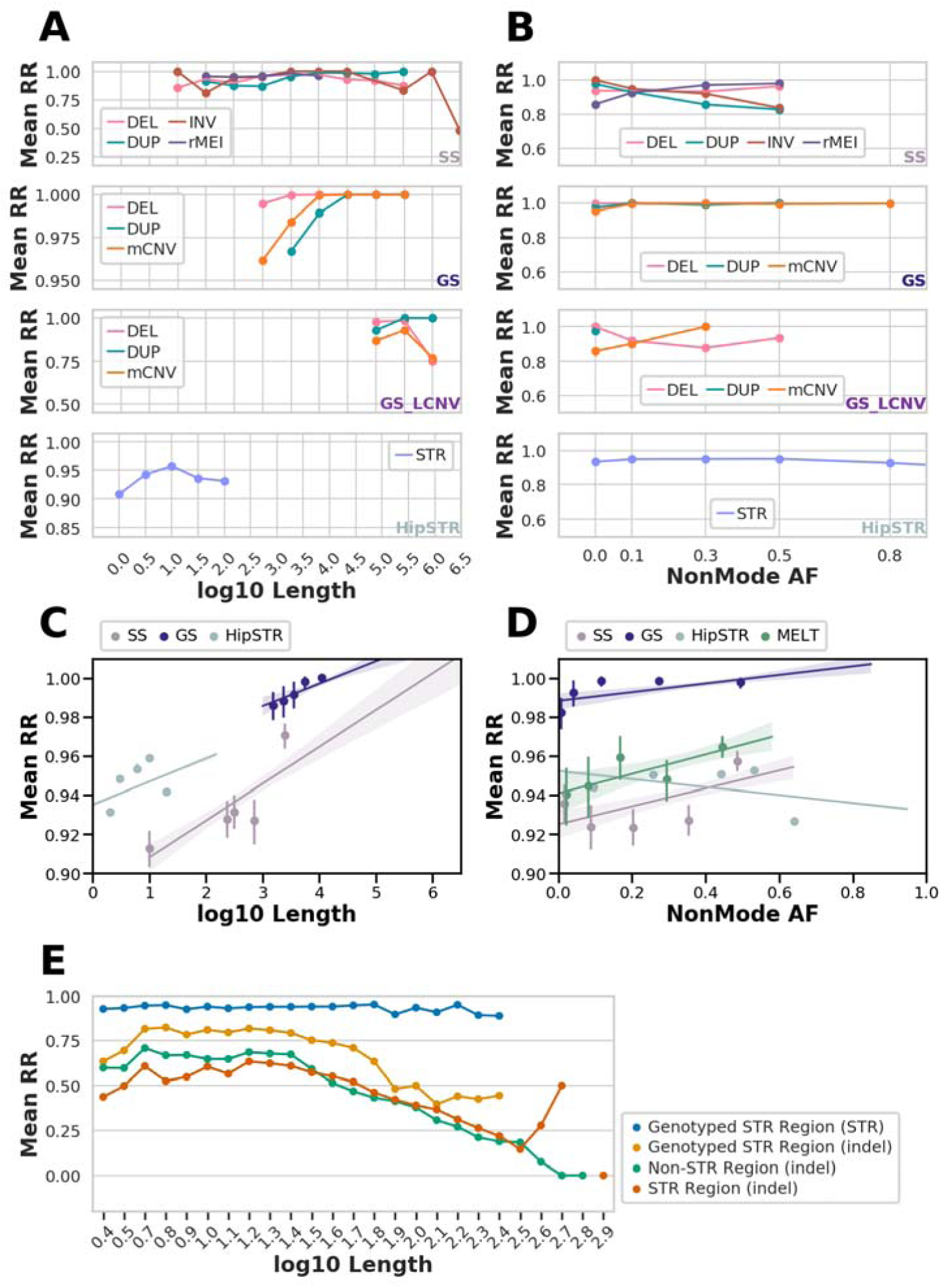
Effect of Length and Allele Frequency on Replication Rate Within Callers. (A and B) Replication rate versus log10 variant length (A) and non-mode allele frequency (B) for deduplicated variants passing filters from each variant caller, stratified by variant class, prior to unifying between variant callers. (C and D) Replication rate versus log10 length for each variant caller after regressing out the effect of non-mode allele frequency (C) and versus non-mode allele frequency (D) after regressing out the effect of log10 length. Points represent the centers of equally sized bins with error bars showing 95% confidence intervals around the mean. Regression lines are shown with shading representing 95% confidence intervals. (E) Replication rate of indels that overlap STR reference regions that are genotyped as non-reference in at least one individual by HipSTR (orange), or are not polymorphic in HipSTR (red), or do not overlap an STR region (green) divided into bins by length. Each point represents the center of a bin.

**Figure S12.**
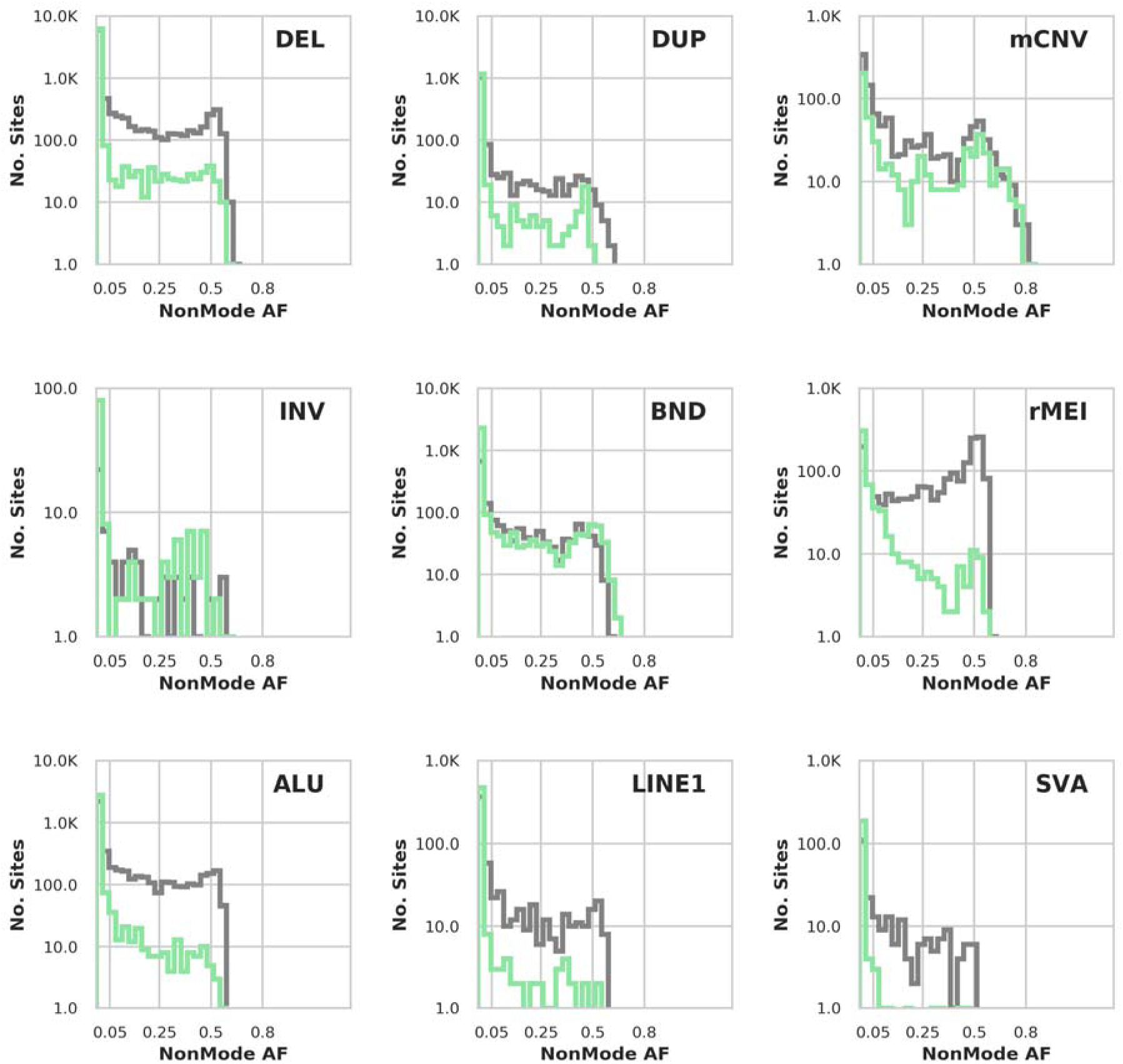
Comparing Allele Frequency Distribution of Known and Novel Variants. Non-mode allele frequency distributions of known (defined as overlapping GTEx or 1KGP variants, gray) and novel variants (defined as not overlapping GTEx or 1KGP variants, green) among unrelated i2QTL samples.

**Figure S13.**
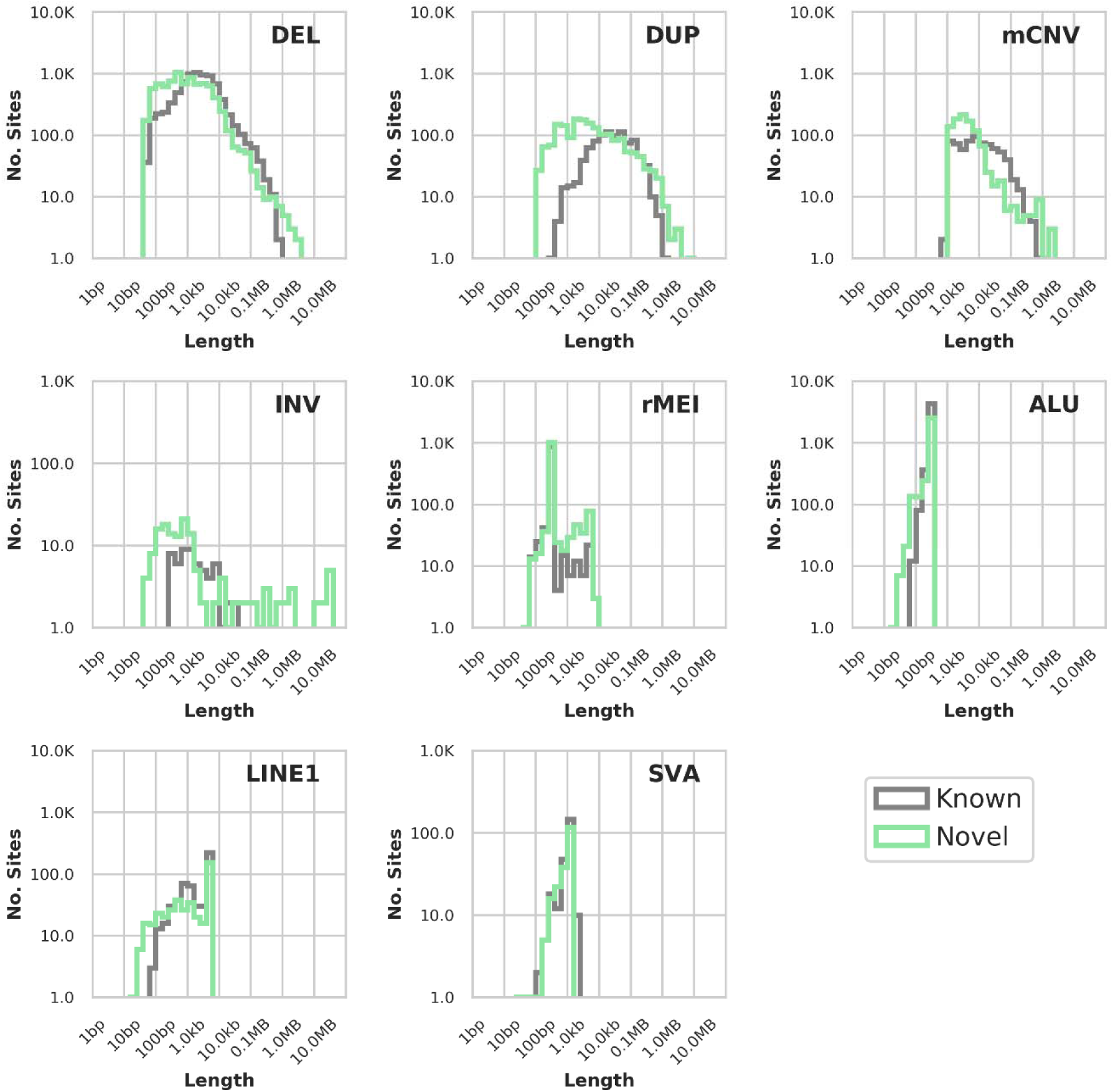
Comparing Length Distribution of Known and Novel Variants. Distribution of variant sizes in known and novel variants among unrelated i2QTL samples after intersection with 1000 Genomes Project and GTEx version 6 SV maps.

**Figure S14.**
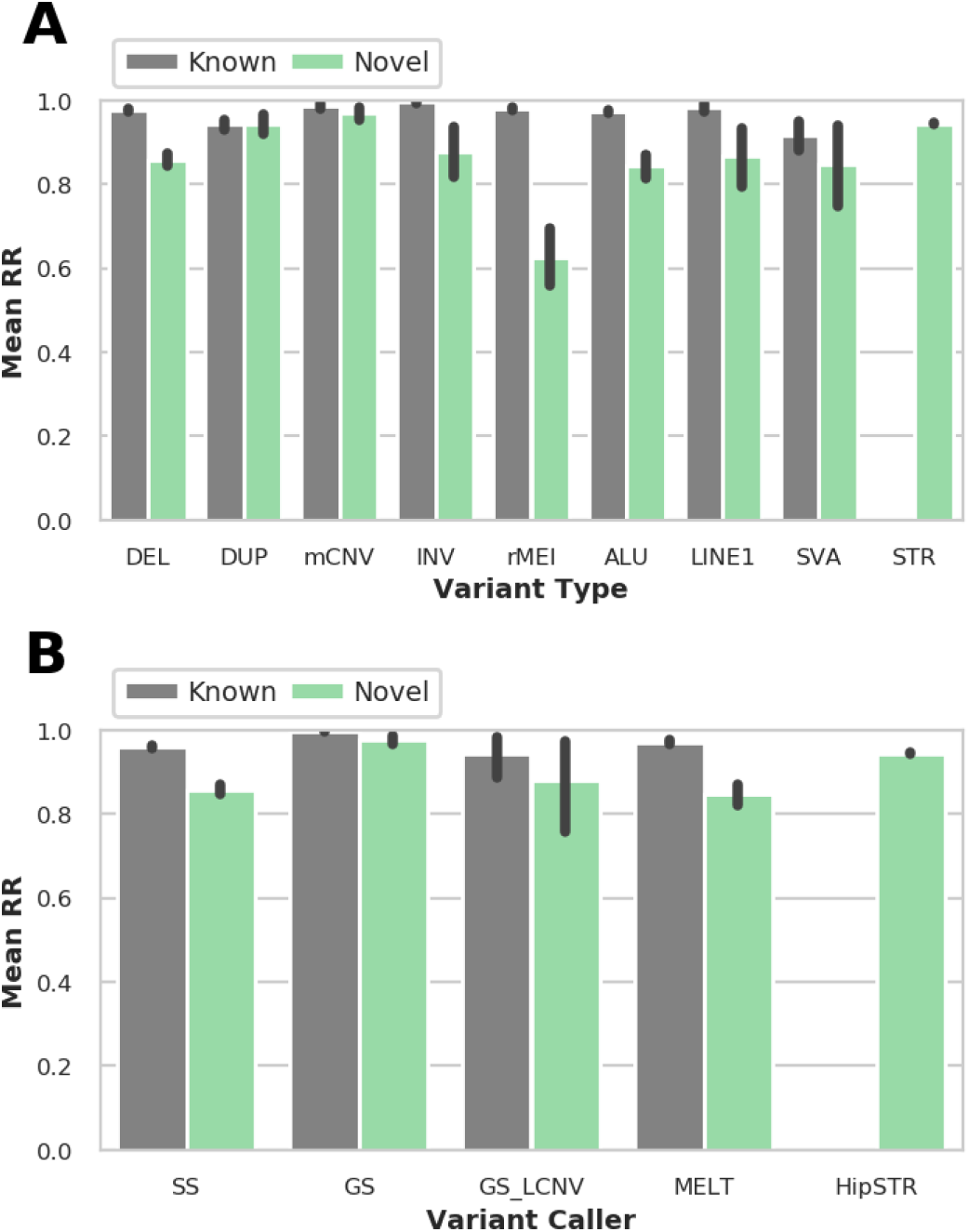
Replication rate of Novel and Known Variants. Average replication rate of novel and known variants (A) by caller and (B) by class. Error bars indicate 95% confidence interval around the mean.

**Figure S15.**
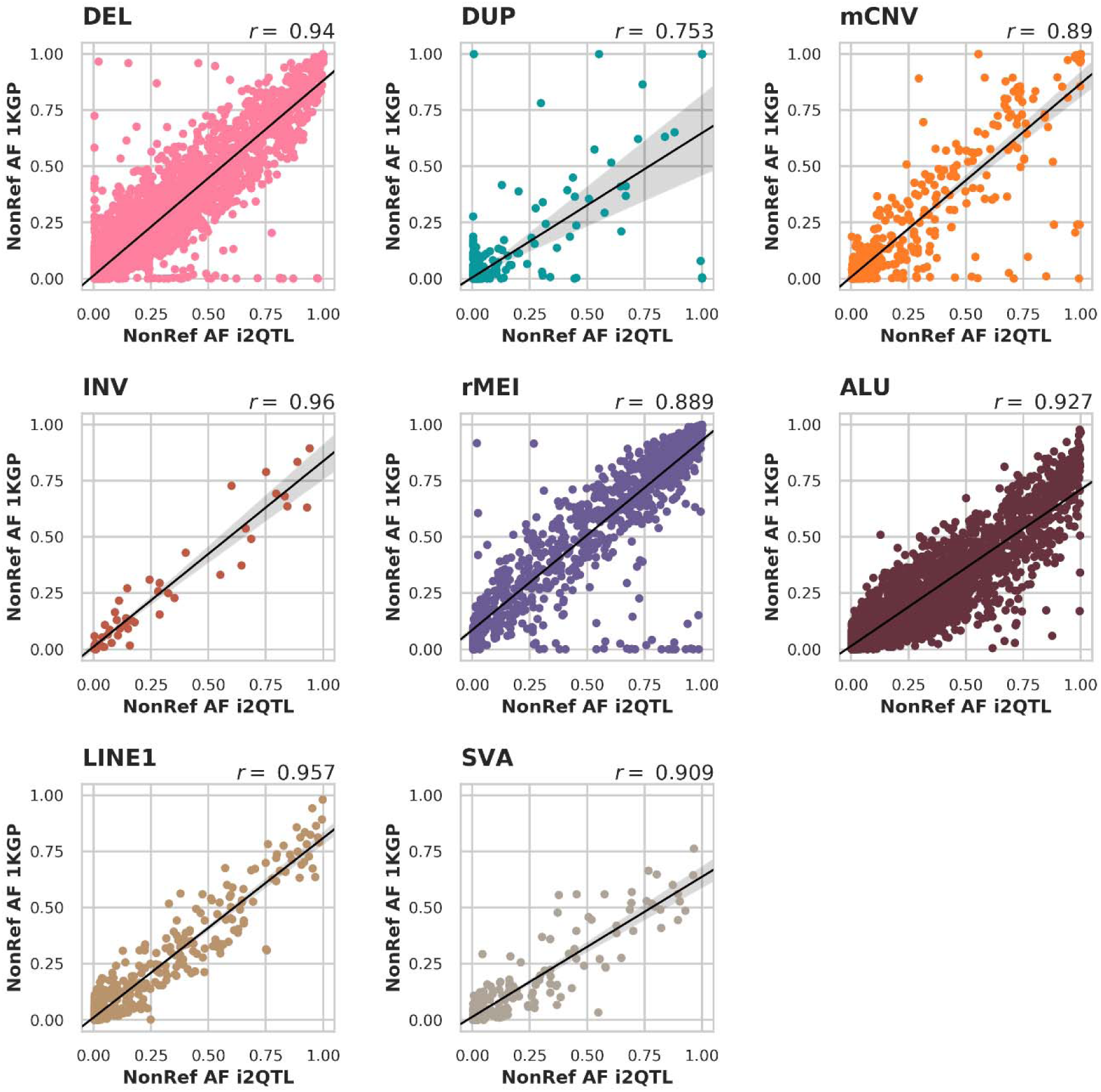
Allele Frequency Comparison of variants co-Discovered in i2QTL and the 1000 Genomes Project. Non-reference allele frequency of overlapping variants in i2QTL and 1KGP(Sudmant et al., 2015), stratified by variant class.

**Figure S16.**
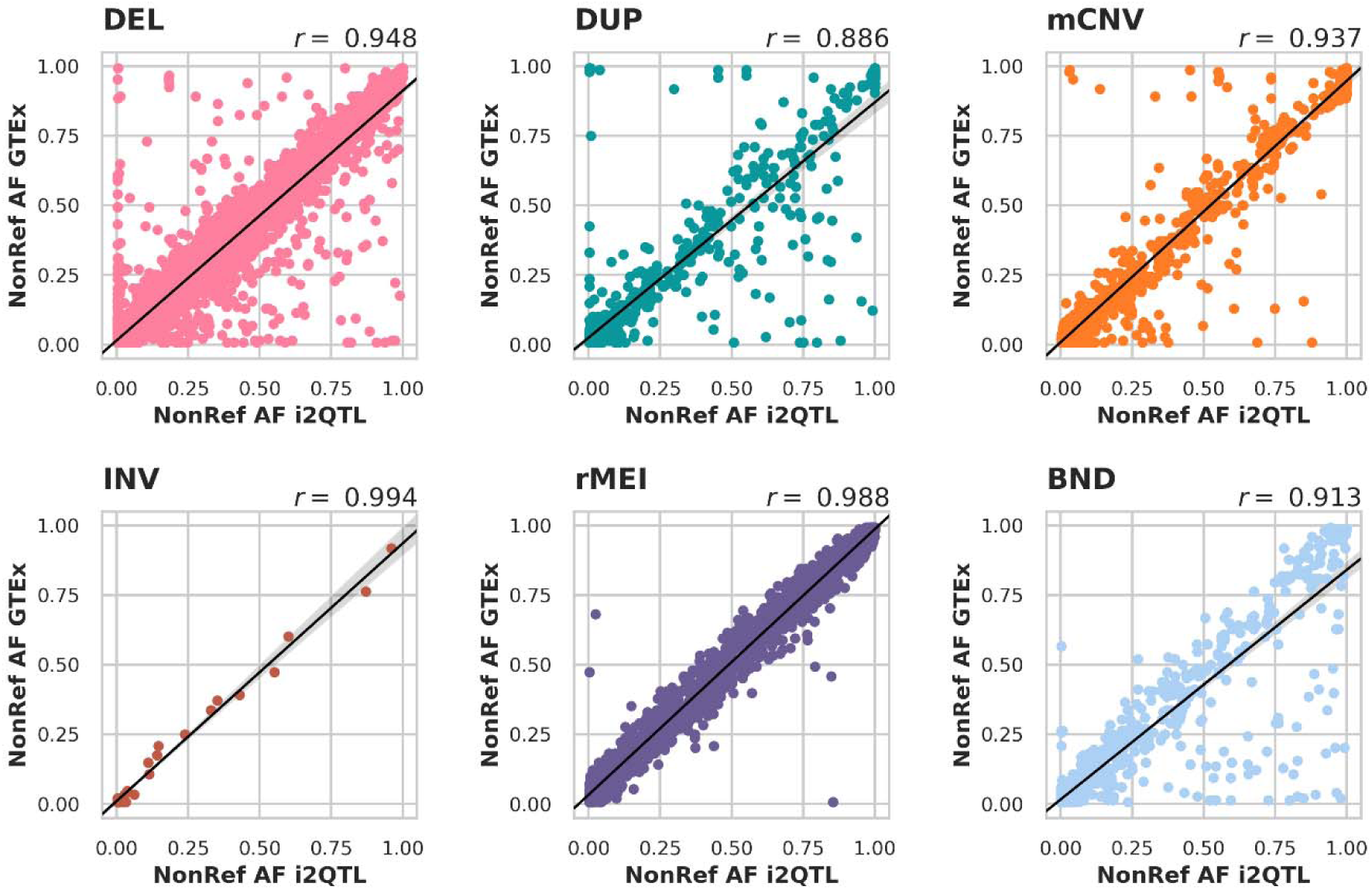
Allele Frequency Comparison of variants co-Discovered in i2QTL and GTEx. Non-reference allele frequency of overlapping variants in i2QTL and the GTEx V.6 SV call set(Chiang et al., 2017) in their respective cohorts, stratified by variant class. For i2QTL variants, the non-reference allele frequency is computed among unrelated samples.

**Figure S17.**
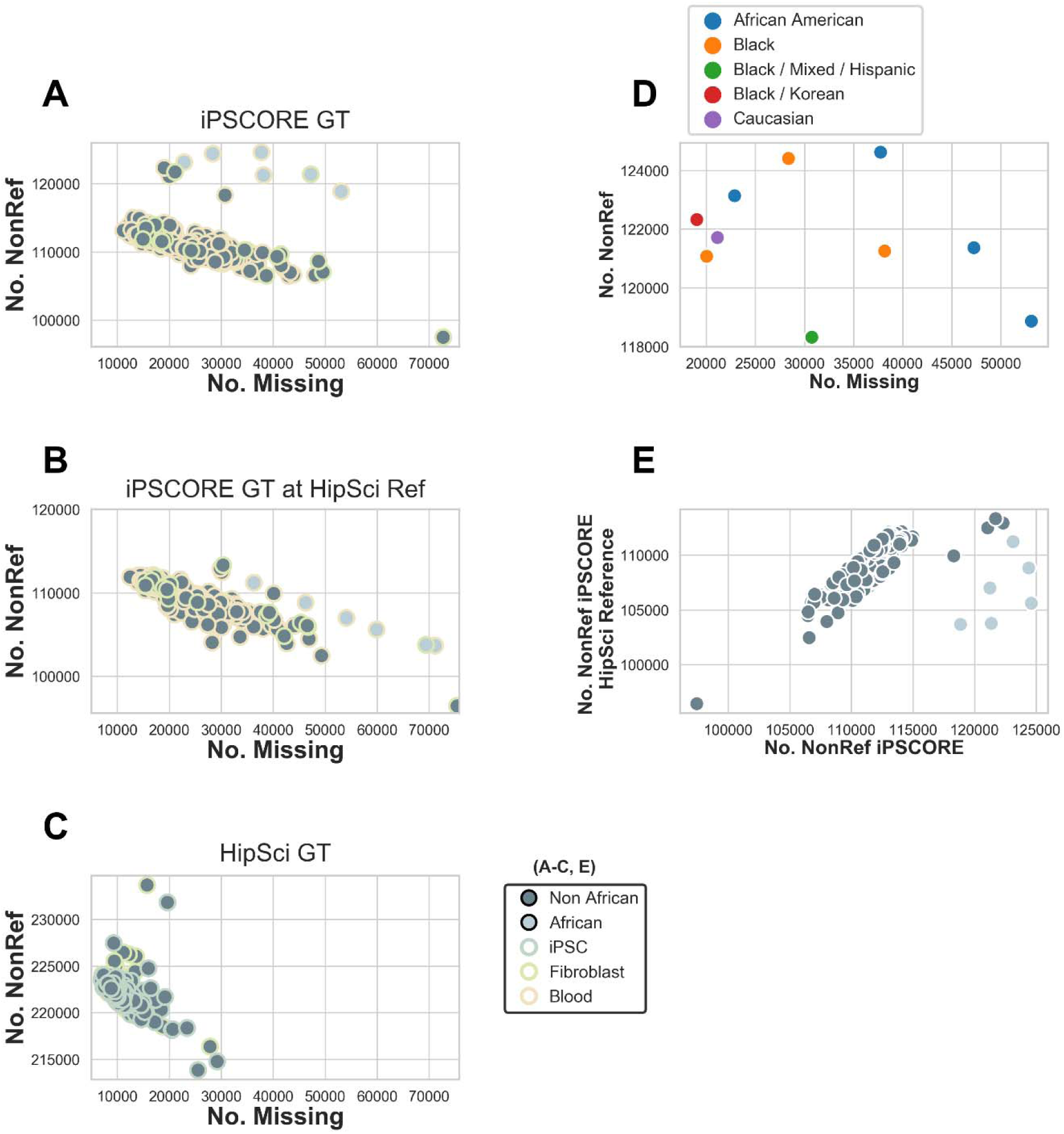
HipSTR Quality Control Before Merging Genotypes from iPSCORE/HipSci. (A-C) Number of non-reference calls per sample for HipSTR genotypes on (A) iPSCORE samples, (B) on iPSCORE samples using HipSci genotypes and (C) on HipSci samples. (D) Outliers from A and B are largely samples from individuals with African predicted super population (shown light grey) or that self-reported as partly African. One iPSCORE outlier sample (A, bottom right) was excluded from call rate filtering (80%) of variants. (E) Number of non-reference genotypes discovered in iPSCORE samples versus number discovered in iPSCORE samples by genotyping HipSci reference alleles. The majority of non-reference sites in iPSCORE were also polymorphic in the HipSci sample set, and the genotypes were similar, however, variants unique to the African samples are not well represented (shown in light grey), as none of the HipSci samples were of African ancestry.

**Figure S18.**
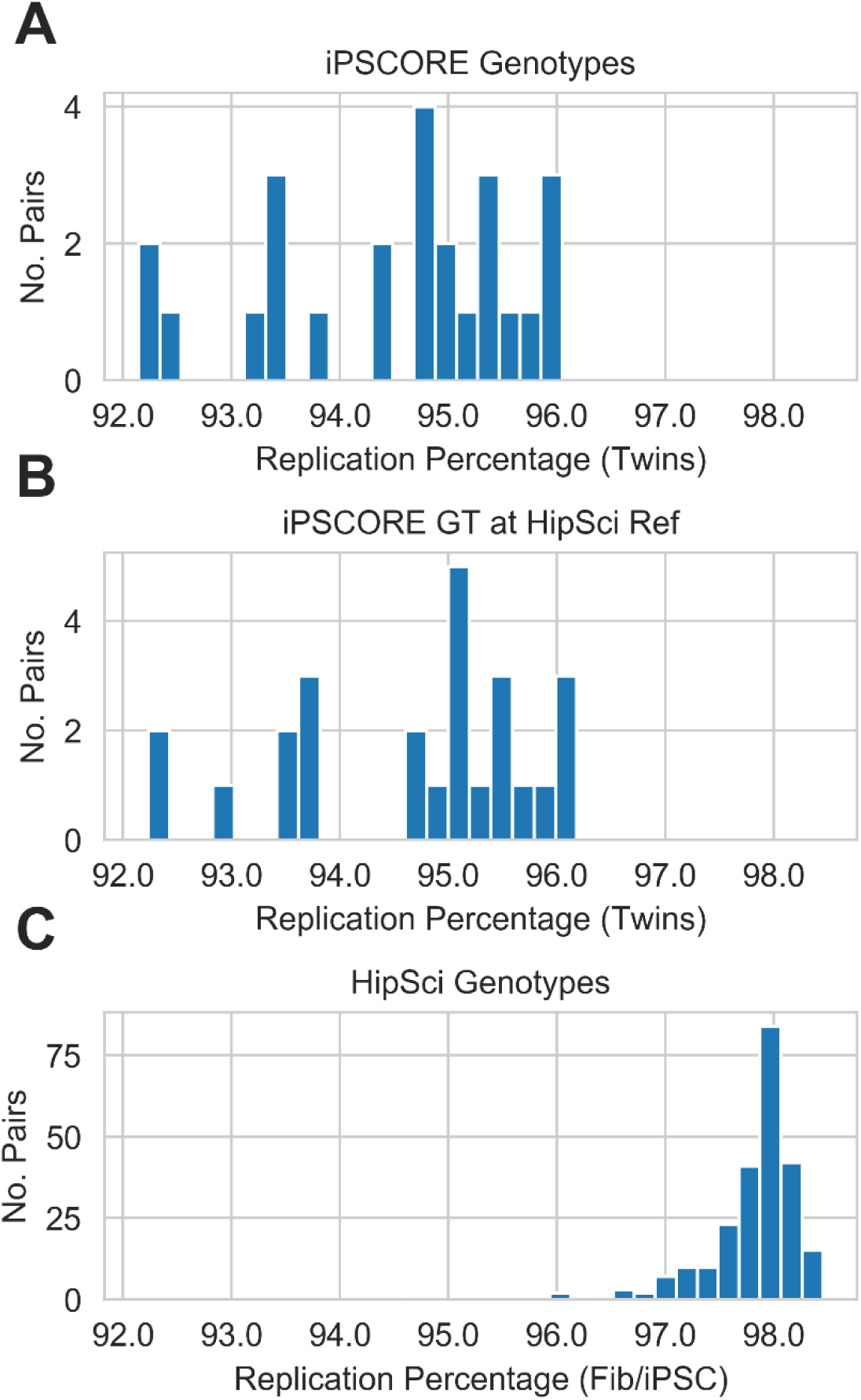
HipSTR Replication Rates in Twin Pairs and iPSC Fibroblast Pairs for Different Genotyping Subsets. (A-C) Replication rate per twin pair after quality filtering for HipSTR genotypes (A) on iPSCORE samples, (B) on iPSCORE samples using HipSci genotypes and (C) on HipSci samples. Here, we observed higher replication percentages among HipSci samples, due to the PCR free protocol of these WGS samples.

